# Structure Primed Embedding on the Transcription Factor Manifold Enables Transparent Model Architectures for Gene Regulatory Network and Latent Activity Inference

**DOI:** 10.1101/2023.02.02.526909

**Authors:** Andreas Tjärnberg, Maggie Beheler-Amass, Christopher A Jackson, Lionel A Christiaen, David Gresham, Richard Bonneau

**Affiliations:** Center for Developmental Genetics, New York University, New York 10003 NY, USA; Center For Genomics and Systems Biology, NYU, New York, NY 10008, USA; Department of Biology, NYU, New York, NY 10008, USA; Flatiron Institute, Center for Computational Biology, Simons Foundation, New York, NY 10010, USA; Courant Institute of Mathematical Sciences, Computer Science Department, New York University, New York, NY 10003, USA; Center For Data Science, NYU, New York, NY 10008, USA; Prescient Design, a Genentech accelerator, New York, NY, 10010, USA; Mortimer B. Zuckerman Mind Brain Behavior Institute, Columbia University, New York, NY, 10010, USA; Sars International Centre for Marine Molecular Biology, University of Bergen, Bergen, Norway; Department of Heart Disease, Haukeland University Hospital, Bergen, Norway

## Abstract

The modeling of gene regulatory networks (GRNs) is limited due to a lack of direct measurements of regulatory features in genome-wide screens. Most GRN inference methods are therefore forced to model relationships between regulatory genes and their targets with expression as a proxy for the upstream independent features, complicating validation and predictions produced by modeling frameworks. Separating covariance and regulatory influence requires aggregation of independent and complementary sets of evidence, such as transcription factor (TF) binding and target gene expression. However, the complete regulatory state of the system, *e.g*. TF activity (TFA) is unknown due to a lack of experimental feasibility, making regulatory relations difficult to infer. Some methods attempt to account for this by modeling TFA as a latent feature, but these models often use linear frameworks that are unable to account for non-linearities such as saturation, TF-TF interactions, and other higher order features. Deep learning frameworks may offer a solution, as they are capable of modeling complex interactions and capturing higher-order latent features. However, these methods often discard central concepts in biological systems modeling, such as sparsity and latent feature interpretability, in favor of increased model complexity. We propose a novel deep learning autoencoder-based framework, *StrUcture Primed Inference of Regulation using latent Factor ACTivity* (SupirFactor), that scales to single cell genomic data and maintains interpretability to perform GRN inference and estimate TFA as a latent feature. We demonstrate that SupirFactor outperforms current leading GRN inference methods, predicts biologically relevant TFA and elucidates functional regulatory pathways through aggregation of TFs.

## 1 Introduction

Transcription factor (TF) regulation of mRNA transcription is a main mechanism through which cells control gene expression and respond to context-specific signals [1, 2]. The relationship between TFs and the genes they control forms an interconnected gene regulatory network (GRN), and interpreting this network is necessary to understand cell and organism heterogeneity, development & differentiation, tissue organization, and diseases [3–6]. GRNs are typically represented as causal graphs, which have regulatory TFs linked to target genes. Regulatory interactions in the GRN are difficult to collectively determine experimentally, so it is necessary to computationally infer the structure of the GRN. This inference is complicated by regulatory relationships between TFs and genes that are context-dependent [7], as TFs may control gene expression differently in different cell types or under different conditions [8].

TFs themselves can be regulated transcriptionally (by changing their mRNA levels), translationally (by changing the amount of protein produced from mRNA), or post-translationally (by modifying the protein to alter localization or DNA binding ability). Transcription factor activity (TFA), the relative ability of a TF to alter the expression of target genes, is the aggregate combination of all TF regulation, and is difficult to measure experimentally. Inferring TFA as a latent model parameter is a core component of several GRN inference methods [9–13]. Generally, TFA is inferred from existing evidence of TF to target gene regulation, combined, using linear models, with the measured expression of the target genes. Although powerful, this framework lacks the flexibility to account for heterogeneity and contextual relations observed in biological systems, and the activity estimates have limited interpretability. Workarounds to contextualize regulatory relationships have been proposed [14, 15], however the inflexibility of the models and the lack of interpretability of latent factors remains an issue.

Using more complex models to better match known transcriptional regulatory biology places numerous demands on optimization and inference machinaries and limits scale; using scalable learning and optimization methods from the deep learning field to meet these needs is an appealing way to infer GRNs. They have been used to model expression and covariance networks [16], to build a sparse representation of gene clusters [17], to group genes into co-regulated modules [18, 19], to do supervised clustering of gene sets [20, 21 and for dimensionality reduction and denoising[22]. However, interpretable deep learning models for gene regulation, which provide biological insight into causal relationships in addition to prediction, have been difficult to construct [23]. Techniques to interpret deep learning latent features often focus on removing latent features to quantify their influence and importance [24]. Feature quantification can use change in mean squared error on removal of input features (COM) [25], backpropagation of the output layer into latent features (Grad-CAM) [26], or forward propagation of the latent layers into the output layer [19]. These techniques cannot be used for model pruning or selection, and are not bounded or informative for model structure evaluation.

Knowledge priming embeds existing evidence into the model structure by constraining connectivity between features. This informs the model of constraints which are not directly measured in the training data. These constraints both improve model selection and strengthen interpretability, as priming with biological interactions allows biological interpretation of the resulting network graph [27]. However, this approach lacks the ability to infer novel regulatory structure and is limited to what has been previously documented, as well as to data sets where all required evidence types are extant. Most target genes are regulated by a limited number of TFs that have no direct effect on other genes, encapsulated in the concept of sparsity in GRNs[28]. Model selection, choosing a limited set of regulators per gene, is therefore a key problem for GRN inference, and many techniques have been applied [29–37]. Deep learning models generally remain overparameterized, making biological interpretation difficult, and techniques must be applied after training to eliminate model parameters and enforce sparsity.

## 2 Results

We present a model inspired by white-box machine learning approaches [38], that we call *StrUcture Primed Inference of Regulation using latent Factor ACTivity* (SupirFactor). This model incorporates knowledge priming by using prior, known regulatory evidence to constrain connectivity between an input gene expression layer and the first latent layer, which is explicitly defined to be TF-specific. This model ties the latent TF layer causally to *informative genes*, and allows this layer to be directly interpreted as transcription factor activity (TFA). This latent layer is then linked to gene expression in an output layer, which is interpreted as an explicitly inferred GRN.

We also adapt new metrics for model interpretation in this context, we define explained relative variance (ERV), a novel concept to interpret the structure of the inferred network graph for any architecture. Briefly, ERV is defined as the change in residual variance when a latent feature is removed from the model, and is used to rank and interpret graph weight importance within the model. Using ERV allows TF to gene interactions to be interpreted through additional latent layers placed between the TF latent layer and the output layer.

Benchmarking across multiple datasets we find that SupirFactor outperforms previous methods using similar frameworks for recovering GRNs. We find that our model uncovers biologically-relevant TFA and predicts biological function of latent aggregates of TFs in deeper layers, suggesting our model is useful for predictive analyses beyond inferring GRNs. In particular we expect to predict activity of specific TFs and to aggregate TFs into regulatory pathways, that we demonstrate on an experimental *S. cerevisiae* data set and a mammalian large single cell PBMC dataset. GRN interpretability and context specific network analysis is facilitated by using ERV and, we demonstrate its utility by applying trained models to context specific and unseen data.

### 2.1 The SupirFactor Regulation Model

The SupirFactor model learns a set of weights connecting genes to TFs in the prior, where these genes functionally serve as reporters for the activity of connected TFs. As opposed to inferring TFA explicitly, as in network component analysis (NCA) (Section 4.4.4) where regulatory evidence is static in the prior, we set *ϕ* = *g*(***W***, ***x***), where ***W*** ∈ ***P*** is the weighted influence of genes to TFs derived from prior evidence ***P***, and ***x*** is the expression of genes informing on ***ϕ*** which may be a subset of *informative genes* connected through ***W***. This extends our model to *x* = *f*(*g*(***W***, ***x***), **Θ**), where **Θ** is the GRN.

We actualise this model using a deep learning framework choosing *f*, and *g* and learn ***W***, and **Θ** with ***ϕ*** as a latent feature prduced by the weighted output from ***x*** mapped through a learned ***W***. Depending on the form of *f*, and including non-linearities, we can learn additional higher order interactions and regulatory pathways (Section 4.1). The complex version of the model that can capture other interactions in *f* we call “hierarchical” SupirFactor (Figure 1A). A simple version of this framework uses a single bottleneck layer for our function *f* we refer to as “shallow” SupirFactor (Figure 1B).

**Figure 1:**
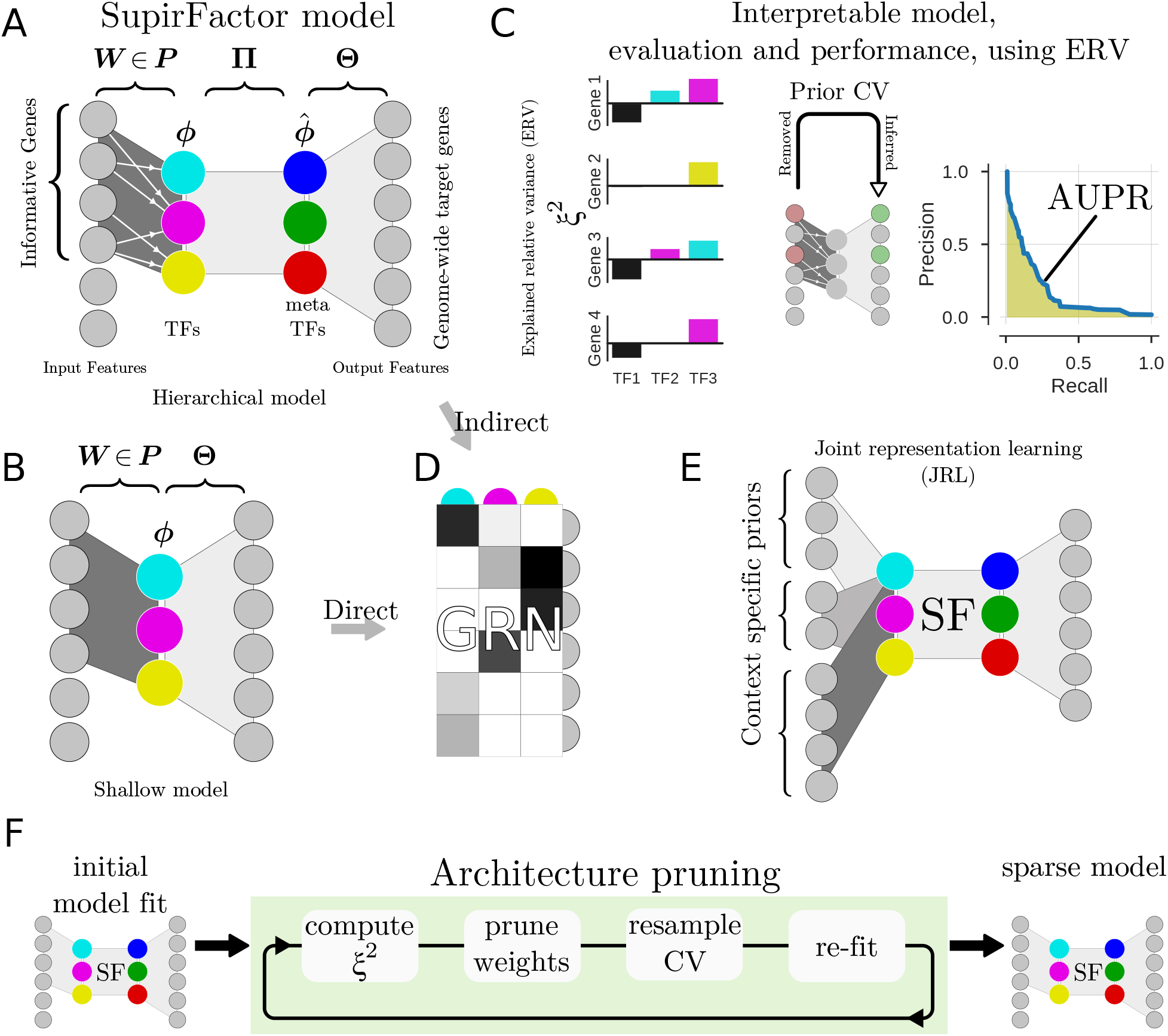
Outline of the SupirFactor framework. The SupirFactor model is constructed like an autoencoder where we embed gene expression data on the transcription factor manifold, exploring two architectures, the “hierarchical” **A**, and the shallow **B** architecture. The output of the first layer defines the latent features marked as TFs (Transcription Factors) and the activation *ϕ* is the transcription factor activity (TFA). The prior ***P*** connect the evidence of TF to a set of informative downstream genes, with learnable weights ***W***. For **A**, **Π** connects the TFs to the latent features, here called the meta TFs (mTFs). **Θ** weights the mTF activity (mTFA) to predict genome wide gene expression profiles. In **B** the TFs directly weights TF to gene influence in **Θ**. **C**: To make the model completely interpretable and transparent we use explained relative variance (ERV) *ξ*^2^. ERV estimate importance of all latent factors influence on model output features. This is then used to evaluate the model and its performance. The GRN is cross validated, where genes to TF connections are held out in the input ***W*** and predicted in the GRN which for the shallow model is *Θ* and for the hierarchical model is 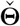 the indirect effect from the TFs to output features. The measured recovery of these links gives insight on stability and biological relevance of the GRN where parameters are ranked by their predictability measured by *ξ*^2^. **D**: Gene regulatory network extracted as indirect TF-gene interaction in hierarchical SupirFactor and direct TF-gene interactions in shallow SupirFactor. **E**: Multi-task learning is implemented in SupirFactor through a joint representation learning (JRL) architecture where biological distinct contexts is independently weighted into a joint GRN representation. **F**: Architecture pruning and sparsity procedure in SupirFactor is used to stabilise and eliminate over-parameterization by eliminating non predicting model parameters facilitated be ERV.

Constructing a prior matrix ***P*** is a challenging but essential task for including informative evidence of regulation. This step is also a way to integrate data types that can shed light on TF-target relationships. This matrix can represent previously known interactions, and it can also encode higher probability interactions derived from chromosome accessibility or TF-chromatin interactions (experimentally measured by ATAC-seq and ChIP-seq) [39, 40]. A more dense ***P*** is likely to include more false positives and will therefore result in an noisier propagation of TF variance. A sparser ***P*** is likely to have many false negatives, limiting the variance that the model is able to explain, resulting in an model that may be less predictive.

A concern is that prior connectivity ***P*** rarely includes reliable sign or weight estimates. Inferring signs for ***P*** from the direction of change after perturbations is technically difficult, as it requires perturbing all TFs included in the model. Relying on expression correlation to infer signs will conflate both indirect and co-varying regulation. We expect that refitting weights ***W*** ∈ ***P*** dependant on **Θ** will mitigate these problems.

### 2.2 Selection of SupirFactor hyperparameters

We evaluate the SupirFactor framework (Figure 1) for GRN inference from bulk and single-cell RNA expression data. First, to test our model setup, explore interpretability, and compare performance to other models, we benchmark using multiple data sets where a partial ground truth network is available (called the gold standard in this work) on the “shallow” SupirFactor. We have previously assembled a GRN inference data package that consists of two prokaryotic *Bacillus subtilis* bulk RNA expression data sets (B1 and B2), two *Saccharomyces cerevisiae* bulk RNA expression data sets (S1 and S2), and one *S. cerevisiae* single-cell RNA expression data set (scY)[41]. This data package also includes a *B. subtilis* gold standard network [10] and a *S. cerevisiae* gold standard network [42], both derived from literature databases. Single-cell RNA expression data is preprocessed (Section 4.7.1) and both bulk and single-cell data is feature normalized (Section 4.7.2) prior to model fitting.

To be able to extract a GRN from our model that gives us latent feature to output connections, we need to be able to interpret model weights, features and their relative importance. This is difficult in multi-layer architectures, and weights may not be scaled with biological importance. Instead, we devise a metric to quantify the direct effect a latent feature has on its targets in the output, Explained Relative Variance (ERV) *ξ*^2^ (Figure 1C), with appealing properties for GRN inference. This metric is based on feature removal [24], and is computed as the coefficient of partial determination (CPD) [43]. We silence each latent model feature and compute the consequent effect on the output variance MSE, scoring the silenced model feature against the full model prediction without retraining (Section 4.4.1).

Deep learning models have a number of hyperparameters that needs to be tuned for optimal model performance. Two critical hyperparameters for GRN inference are the *weight-decay*, typically called the L2 penalty, and dropout, stochastic perturbation of the data during training to attenuate noise and improve model generalisation (Section 4.3). We test dependence on these hyperparameters by searching for L2 and dropout (on input and latent features) with the simplified shallow SupirFactor model. Each hyperparameter value is tested by splitting expression data 50-50 into training and validation set, using 80% of the gold standard network as prior network information for the model and holding out 20% for scoring. Negative controls consist of either shuffling the data or the prior network. Area under precision recall curve (AUPR) is used to score the network structure against the gold standard network, and *R^2^* is used to quantify prediction accuracy. Each configuration is rerun repeatedly and average performance is reported (Supplemental Figure S2–S9).

We observe that in some cases (for S2 and B1 datasets), when increasing L2 beyond a specific value, *R*^2^ decreases while AUPR increases (Supplemental Figure S3 & S4). We interpret this as overfitting to the prior network structure and increasing recovery of the gold standard in cases where the these two structures align, such as in scale-free networks with dense highly connected TFs. The model then emphasises these TFs at the cost of prediction accuracy and inclusion of less connected TFs. This is undesirable, so we select hyperparameters that maximises *R*^2^ while maintaining a high AUPR. Negative controls perform as expected; shuffling the data eliminates the biological interpretability and predictive power of the resulting GRN, and shuffling the prior network eliminates only biological interpretability while still achieving good predictive power.

To determine an optimal L2 we look for where *R*^2^ is maximized. We find that prediction accuracy is maximised in the span *R*^2^ ∈ 10^−6^ − 10^4^ for all data sets, where we select an L2 of 10^−4^ for B1 and B2 and 10^−6^ for S1, S2 and scY for further comparisons (Supplemental Figure S2C–S9C). For dropout, in general, we find that setting larger dropout on input and smaller dropout on latent features increases AUPR while maintaining a higher *R*^2^.

### 2.3 SupirFactor benchmarking demonstrates improved biological regulatory network recovery

We summarize the performance of the shallow SupirFactor model using a linear activation function with several modeling choices (Figure 2A). Using ERV as an estimator for biological relevance outperforms interpretation based on model weights alone, as determined by AUPR. Selecting the optimal dropout hyperparameters based on maximum *R*^2^ for the selected L2 (max R2) improves model *R*^2^ at a cost of decreasing network prediction performance (AUPR), when compared to setting a fixed dropout (fixed; input=0.5 and latent=0).

**Figure 2:**
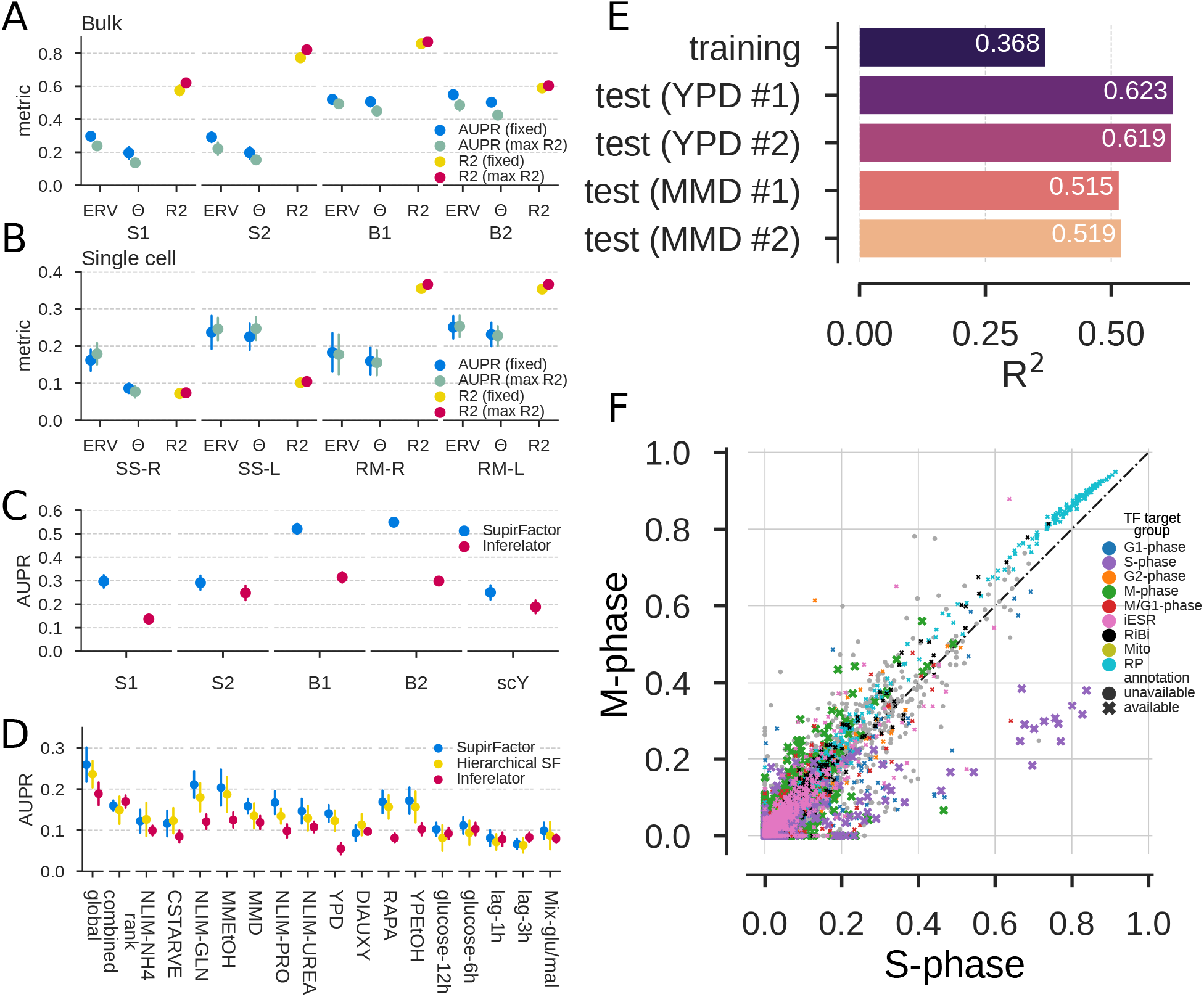
SupirFactor benchmark and hyperparameter evaluation. Performance is evaluated on gold-standard networks consisting only of edges held out of the prior network ***P***, measuring recovery using AUPR. *R^2^* is computed on a validation set of 50% of the data samples held out of the training data. Data sets are labeled for species, with *B. subtilis* (B1 and B2), *S. cerevisiae* bulk RNA expression (S1 and S2), and *S. cerevisiae* single-cell RNA expression (scY). **A**: Comparing network interpretation using model weights (Θ) to interpretation using Explained Relative Variance (ERV), measured using AUPR against edges held out of the prior network ***P***. *R*^2^ is calculated from the full network model. Model hyperparameters are set based on Supplemental Figure S2–S9 and Section 4.3. **B**: Comparing normalization and activation functions for single-cell RNA expression data, as in (**A**). SS- is normalizing to mean of zero and unit variance, RM- is normalizing to maintain a minimum value of 0 (retaining sparsity), -L is linear activation, and -R is ReLU activation. **C**: Benchmarking SupirFactor with optimal parameters selected from (**B**) against a comparable GRN inference method, the Inferelator. **D**: Comparing multi-context network performance between shallow SupirFactor, Hierarchical SupirFactor, and the multi-task Inferelator. GRNs are learned from single-cell (scY) data, with context/task groupings determined by growth condition. Global GRNs are learned from the data without separate groupings (using StARS-LASSO for the Inferelator [41]). Context networks are computed post-training in SupirFactor and split here on growth condition. **E**: Evaluating model prediction *R*^2^ on four novel test data sets, using a GRN trained by Hierarchical SupirFactor. **F**: Comparing contextual network for GRNs defining cell cycle M-phase and S-phase (Table 1). Each point is an interaction from the two contextual networks, colored by the target gene functional annotation. X and Y axis are *ξ*^2^ of the S-phase and M-phase networks. GRN interactions targeting S-phase genes (purple) have higher ERV in the S-phase contextual network, and interactions targeting M-phase genes (green) have higher ERV in the M-phase contextual network.

For single-cell RNA expression data (Figure 2B) we extend the comparisons, comparing linear and rectified linear (ReLU) activation functions (Section 4.8), and comparing standard normalization to a robust sparsity-retaining normalization (Section 4.7.2). RobustMinScaler outperforms StandardScaler normalisation, implying that preserving sparse data for model training is advantageous (Figure 2B). To summarize these benchmarking results, ERV should be used to evaluate model parameters, and dropout hyperparameters can be fixed without loss of prediction accuracy and GRN recovery.

Finally, we compare shallow SupirFactor performance to that of the Inferelator (Figure 2C), a method which takes comparable RNA expression and prior network inputs and learns a GRN. We note that other methods

**Table 1:**
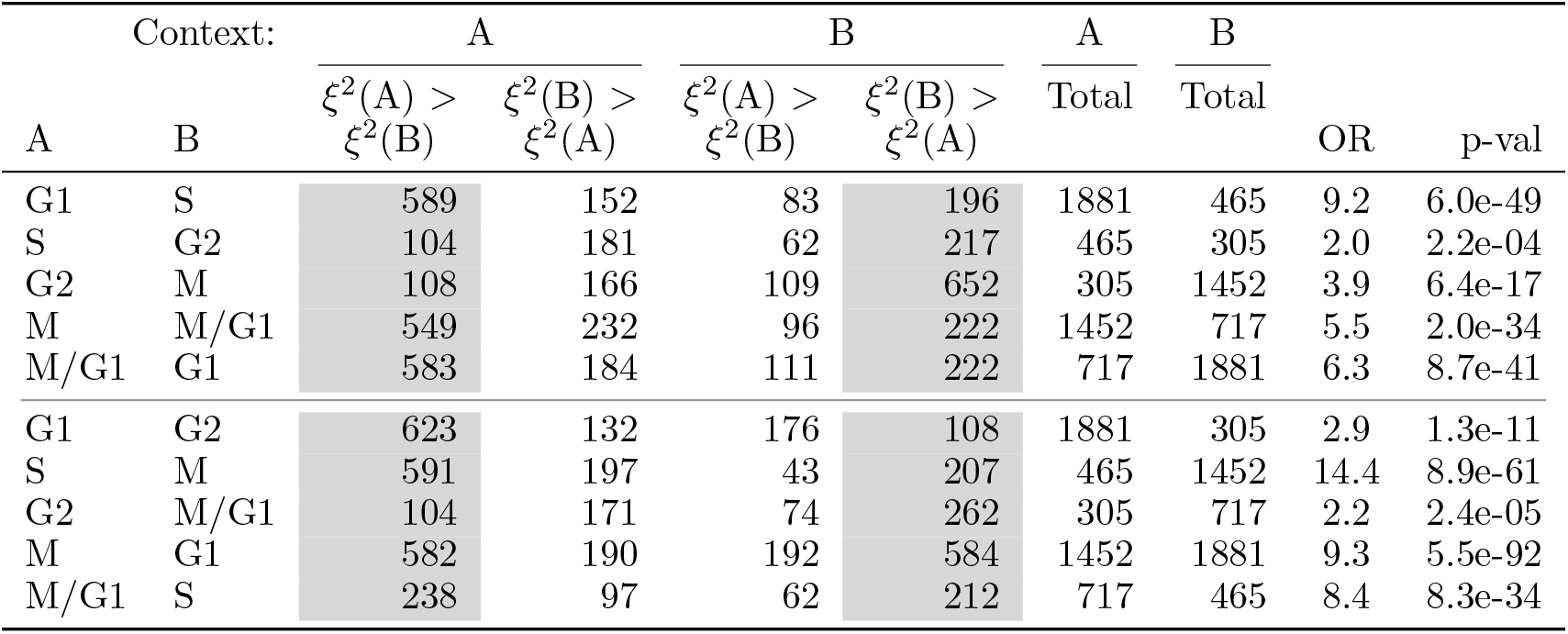
Pairwise comparisons between context-specific cell cycle phase networks from hierarchical Supir-Factor. Network edges are classified into cell cycle phase context based on the existing annotation of the target gene. ERV of the network edges (where *ξ*^2^ > 0.01 in one or more contexts) is then compared as a contingency table. Network edge counts in gray columns are edges where ERV is higher in the correct context-specific network, based on existing annotations. The total column counts network edges (where *ξ*^2^ > 0 for that context) for the context-specific network, and does not total the columns of the contingency table. Odds Ratio (OR) and p-values are calculated by one-sided fisher exact test, using the contingency table.

| Context: |  | A |  | B |  | A | B | OR | p-val |
| --- | --- | --- | --- | --- | --- | --- | --- | --- | --- |
| A | B | $\xi^2(A) > \xi^2(B)$ | $\xi^2(B) > \xi^2(A)$ | $\xi^2(A) > \xi^2(B)$ | $\xi^2(B) > \xi^2(A)$ | Total | Total | | |
| G1 | S | 589 | 152 | 83 | 196 | 1881 | 465 | 9.2 | 6.0e-49 |
| S | G2 | 104 | 181 | 62 | 217 | 465 | 305 | 2.0 | 2.2e-04 |
| G2 | M | 108 | 166 | 109 | 652 | 305 | 1452 | 3.9 | 6.4e-17 |
| M | M/G1 | 549 | 232 | 96 | 222 | 1452 | 717 | 5.5 | 2.0e-34 |
| M/G1 | G1 | 583 | 184 | 111 | 222 | 717 | 1881 | 6.3 | 8.7e-41 |
| G1 | G2 | 623 | 132 | 176 | 108 | 1881 | 305 | 2.9 | 1.3e-11 |
| S | M | 591 | 197 | 43 | 207 | 465 | 1452 | 14.4 | 8.9e-61 |
| G2 | M/G1 | 104 | 171 | 74 | 262 | 305 | 717 | 2.2 | 2.4e-05 |
| M | G1 | 582 | 190 | 192 | 584 | 1452 | 1881 | 9.3 | 5.5e-92 |
| M/G1 | S | 238 | 97 | 62 | 212 | 717 | 465 | 8.4 | 8.3e-34 |

which take these inputs are not amenable to scoring on holdouts that we are employing for our benchmark [41], making comparison difficult. Using the configurations from our benchmarking, SupirFactor improves significantly over previous results on all of the data sets tested. Comparison of the Inferelator to several alternate methods on the same data sets used here has been previously shown [41].

### 2.4 Hierarchical SupirFactor with non-linear activation facilitates interpretation of latent feature activation as biological activity in cell contexts

To account for more complex, multi-TF regulatory interactions we extend our mode in multiple ways, introducing non-linearities and adding additional latent layers that represent interactions. Non-linear functional forms are necessary in the proposed hierarchical SupirFactor architecture to model interactions more complex than linear relationships. This is necessary as GRNs are context- and cell-type-dependent; and thus a TF to gene regulatory interaction may *e.g*. exist when the organism is in one state, but be inaccessible in another.

SupirFactor can distinguish contextual networks by embedding context specific assigned data and computing ERV only within that data set. We explore learning context-dependent GRNs here; we evaluate this ability on the single-cell RNA expression data set, which has samples annotated by growth conditions (Figure S1A). We compare hierarchical SupirFactor (Figure 1A), shallow SupirFactor (Figure 1B), and a comparable multi-task learning approach (AMuSR) in the Inferelator that also learns context-dependent networks. Both the shallow SupirFactor and hierarchical SupirFactor outperform the Inferelator (Figure 2D). Shallow Supir-Factor outperforms the hierarchical SupirFactor in some contexts, although the shallow model uses a linear activation function, and the hierarchical model uses a ReLU activation function. As this activation function constrains the latent features to be strictly positive, latent features are interpretable in the hierarchical model.

We use hierarchical SupirFactor to construct context-specific GRNs for cell cycle phases, by inferring cell cycle phase from transcriptional markers (Figure S1B). Regulatory edges that are actively used in a context-specific GRN should explain more relative variance, compared to edges which are inactive. For each cell cycle we used the gene annotation of cell cycle phase genes and compare ERV between phases for each relevant gene (Table 1). We compare both neighbouring phases and phases skipping the immediate following phase. Computing a one sided fisher exact test classifying genes in the GRN where they have the highest ERV, only using ERV that has *ξ*^2^ > 0.01 in at least 1 of the conditions, we find that for all comparisons we have a strong enrichment of the phases relevant genes in terms of ERV in the corresponding phases.

Hierarchical SupirFactor introduces a larger set of learnable weights and a potential over-parameterisation of the model, and, thus, the expanded model presents new model selection challenges in the context of sparse GRN inference. We test if an iterative method for removing model weights could be used without loss of model performance. ERV is used to rank model weights, and an *ξ*^2^ ≤ 0 indicates a non predictive model weight. Identifying these non-predictive links, pruning them, and then refitting the model attenuates over-parameterization (Figure 1E). After 1 iteration of pruning, nonzero weights reduce to ~ 57% ∈ **Θ**, 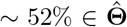 and ~ 80% ∈ **Π** of the pre-pruning model trained on scY. It is critical to determine if models learned by hierarchical SupirFactor generalize. We evaluate the predictive ability of a hierarchical SupirFactor model trained on data set scY using prior knowledge ***P*** without holdouts (Figure 2B, RM-R). To do this, we generate four new experimental test single cell RNA expression data sets by collecting and sequencing cells grown in environmental conditions seen in the training data set (YPD and MMD). This model explains the variance of the training data well (*R*^2^ = 0.37), and also explains the variance of the YPD (*R*^2^ = 0.62) and MMD test data sets (*R*^2^ = 0.52) well (Figure 2E). We conclude that SupirFactor generalises and predicts expression patterns of new data even when model weights have been removed.

The SupirFactor model explicitly fits an intermediate layer which can be directly interpreted as latent Transcription Factor Activity (TFA) for each TF, when this layer uses a ReLU activation function [44]. Using hierarchical SupirFactor, we calculate latent TFA for all TFs. When examining the role of cell cycle TFs, the advantages of TFA are apparent. The TFA for cell cycle TFs is maximal in phases the TFs are expected to regulate (Figure 3A), based on known TF roles from literature[45]. Almost all cells have non-zero TFA for cell cycle TFs in at least one phase of the cell cycle, but TF expression is highly sparse, complicating causal linkage to targets based on TF expression (Figure 3B). We note that for these cell cycle TFs, the expression of the TF often peaks in the phase before the TFA of the TF.

**Figure 3:**
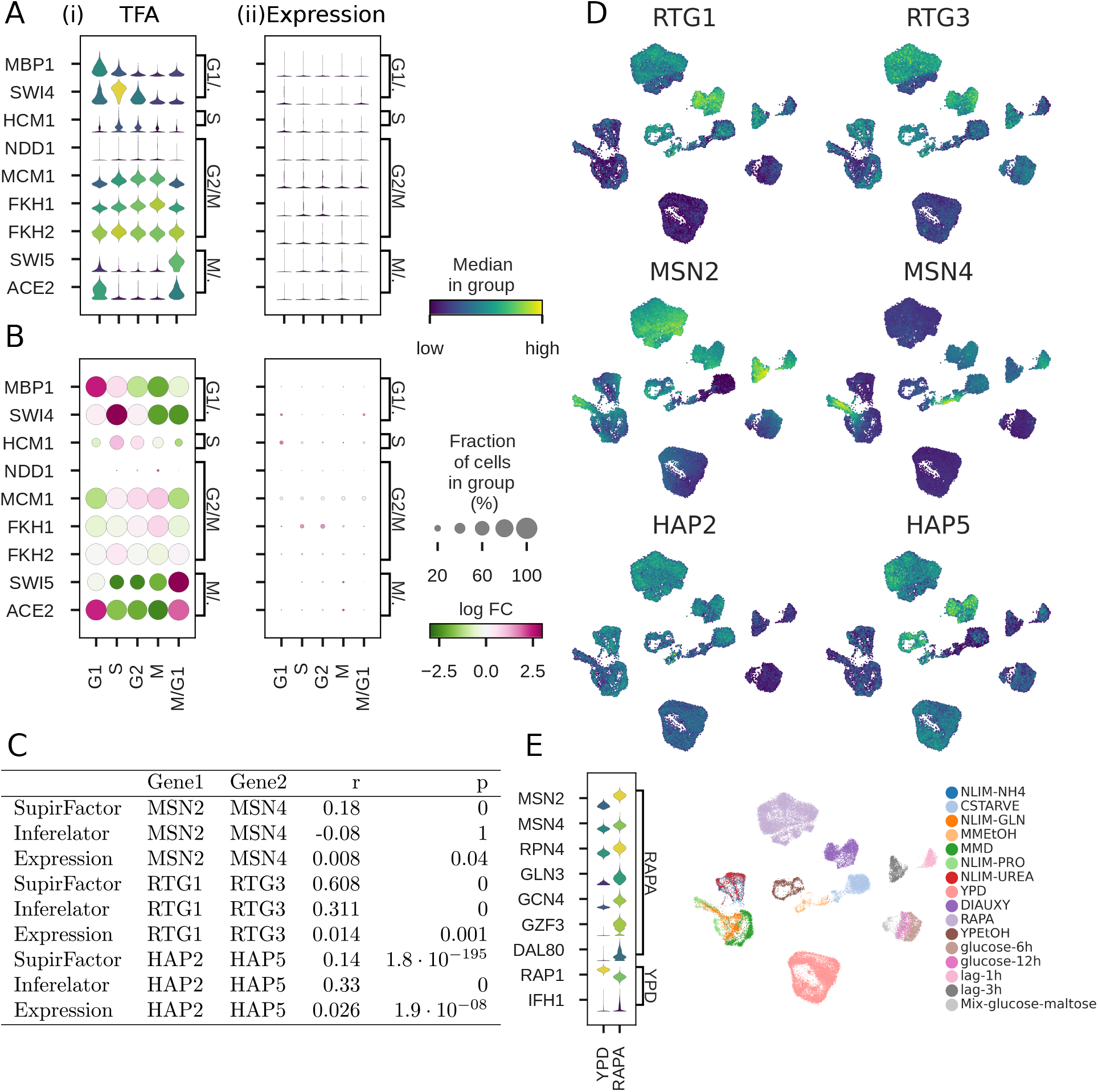
Transcription factor activity in single cell yeast. TFA estimated from hierarchical SupirFactor model. Violin plots are generated by scaling [0, 1] the underlying measurement. **A**: Cell cycle TFs regulate gene expression in specific cell cycle phases, with the phase the TF regulates annotated on the right y-axis [45]. Cell cycle phase of each cell is inferred from transcriptome, and annotated on the x-axis. Panel (i) plots TFA, and panel (ii) plots the RNA expression of the TF. **B**: Same as **A**. Dot size represents the percentage of cells with a non-zero value. Color represents log fold-change (log FC) across the cell cycle phases. Panel (i) plots TFA, and panel (ii) plots the RNA expression of the TF. **C**: Interacting TF pair pearson correlation for SupirFactor TFA, Inferelator TFA, and TF expression **D**: Comparing TFA between rapamyacin (RAPA) treated and untreated (YPD) cells TFs known to be activated by treatment [46] or known to be more active in untreated cells are annotated on the right y-axis. **E**: UMAP projection of the scY dataset showing TFA estimate of co-regulator TFs and growth conditions.

We also wanted to test the role of functionally related TFs. Transcription factors often interact with each other to define regulatory states, as part of multisubunit complexes, or by competing for the same DNA binding regions. These interactions should be reflected in how TF activity correlates, even if these interactions are not explicitly embedded in the model. We select several typical examples of TF pairs with known interactions and compare inferred TFA between hierarchical SupirFactor and the Inferelator (Figure 3C). MSN2 and MSN4 are partially redundant stress response TFs that bind to the same DNA motif, and we expect their activity to be partially correlated. Hierarchical SupirFactor TFA is weakly correlated (*r* = 0.18), unlike expression of MSN2 and MSN4, which are uncorrelated (*r* = 0.01). RTG1 and RTG3 are obligated to form a physical dimer for functionality, and we expect their activity to be strongly correlated. Hierarchical SupirFactor TFA is strongly correlated (*r* = 0.61), unlike expression of RTG1 and RTG3, which are uncorrelated (*r* = 0.01). Finally, HAP2 and HAP5 are part of the multisubunit heme-activated TF complex, and we expect their activity to be strongly correlated. In this case, hierarchical SupirFactor is less successful at correlating TFA (*r* = 0.14) than the Inferelator (*r* = 0.33), expression is again uncorrelated (*r* = 0.03). Overlaying TFA onto a reduced-dimensionality plot allows for the comparison between TF activities and the experimental conditions which cause them to be correlated or uncorrelated (Figure 3D).

Finally, we compare the TFA between perturbed, rapamycin-treated cells and untreated control cells (Figure 3E). Rapamycin is expected to inhibit TOR pathway signalling, altering stress response and nutrient response TF activities [45]. By comparing the TFA between perturbed and control cells, hierarchical SupirFactor is able to reconstruct which TFs are activated and deactivated by this perturbation.

### 2.5 Hierarchical SupirFactor combines TFs into pathways

In hierarchical SupirFactor we introduce an additional latent layer, which we interpret as meta transcription factors (mTFs) that aggregate TFs into multi-regulator pathways. As this mTF layer is directly connected to the output gene expression, we expect that the mTF layer activity (mTFA) can be interpreted as the activity of a regulatory pathway.

To test this hypothesis, we explore the hierarchical SupirFactor model trained on the single-cell yeast data (scY). mTF functions are determined by enrichment for regulation of genes that are annotated with Kyoto Encyclopedia of Genes and Genomes (KEGG) pathways [47]. As this data was collected from cells growing in different carbon and nitrogen sources, we focus on enrichment of a specific subset of metabolic pathways. 42 of 148 mTFs are enriched for target genes (defined as mTF to gene connections with *ξ*^2^ > 0.1) to these metabolic KEGG pathways (Figure 4A).

**Figure 4:**
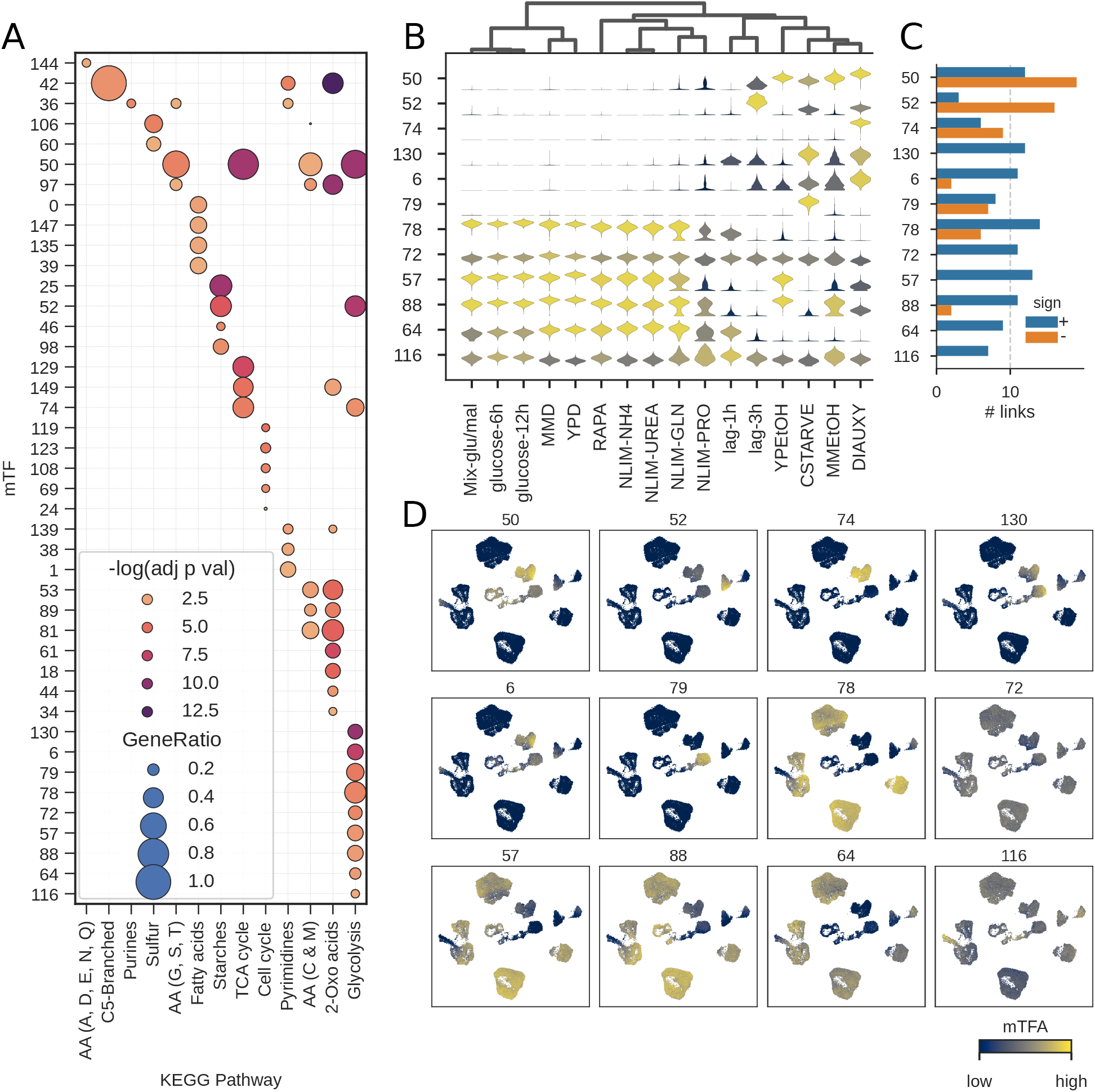
Meta transcription factor (mTF) functional enrichment analysis in single cell yeast. TFA estimated from hierarchical SupirFactor model trained in Figure 3. mTFs are nodes in the SupirFactor model Φ latent layer and are numbered from 1-148. **A** Pathway enrichment of mTFs to selected core metabolic KEGG pathway annotations on target genes. **B** mTF activity for cells in each growth condition (mTF activity scaled [0, 1] for comparison). **C** Positive (activating) and negative (repressing) weights from mTFs to target genes within the Glycolysis KEGG-pathway for each mTF. **D** mTF activity for cells overlaid on a low-diensional UMAP projection. Cell metadata plotted in Supplemental Figure S1.

We focus on the 12 mTFs which are enriched for glycolysis target genes, a pathway which is a core part of the central carbon metabolism. Many of the growth conditions in the training single-cell yeast gene expression data set use glucose as a primary carbon source (Supplemental Figure S1E), and we see that these conditions have very similar mTF activation (Figure 4B). The remaining growth conditions use different carbon sources, requiring different regulation of the central carbon metabolism. Six of the glycolysis mTFs are activated in these non-glucose carbon sources, but have considerably more repressive (negative weighted) links to target genes (Figure 4C), suggesting that they are mainly downregulating glycolytic genes. We can further overlay mTF activity onto a low-dimensionality projection in order to identify mTFs which are linked to carbon source with little heterogeneity (*e.g*. 74 & 79) and which have heterogeneity within growth conditions (*e.g*. 57 & 78) (Figure 4D). We observe that mTFs aggregate biologically functional groups in their targets and can be evaluated quantitatively as activities of these pathways.

### 2.6 SupirFactor models regulation of mammalian PBMCs with multimodal single cell sequencing data

We evaluate the use of SupirFactor to model complex biological systems by applying the method to model Peripheral Blood Mononuclear Cell (PBMC) gene regulation, using a paired multi-omic single cell ATAC-seq and RNA-seq dataset [48] (Figure 5A). These two data types are integrated by using ATAC-seq chromatin accessibility as a cell-specific mask (Section 16).

**Figure 5:**
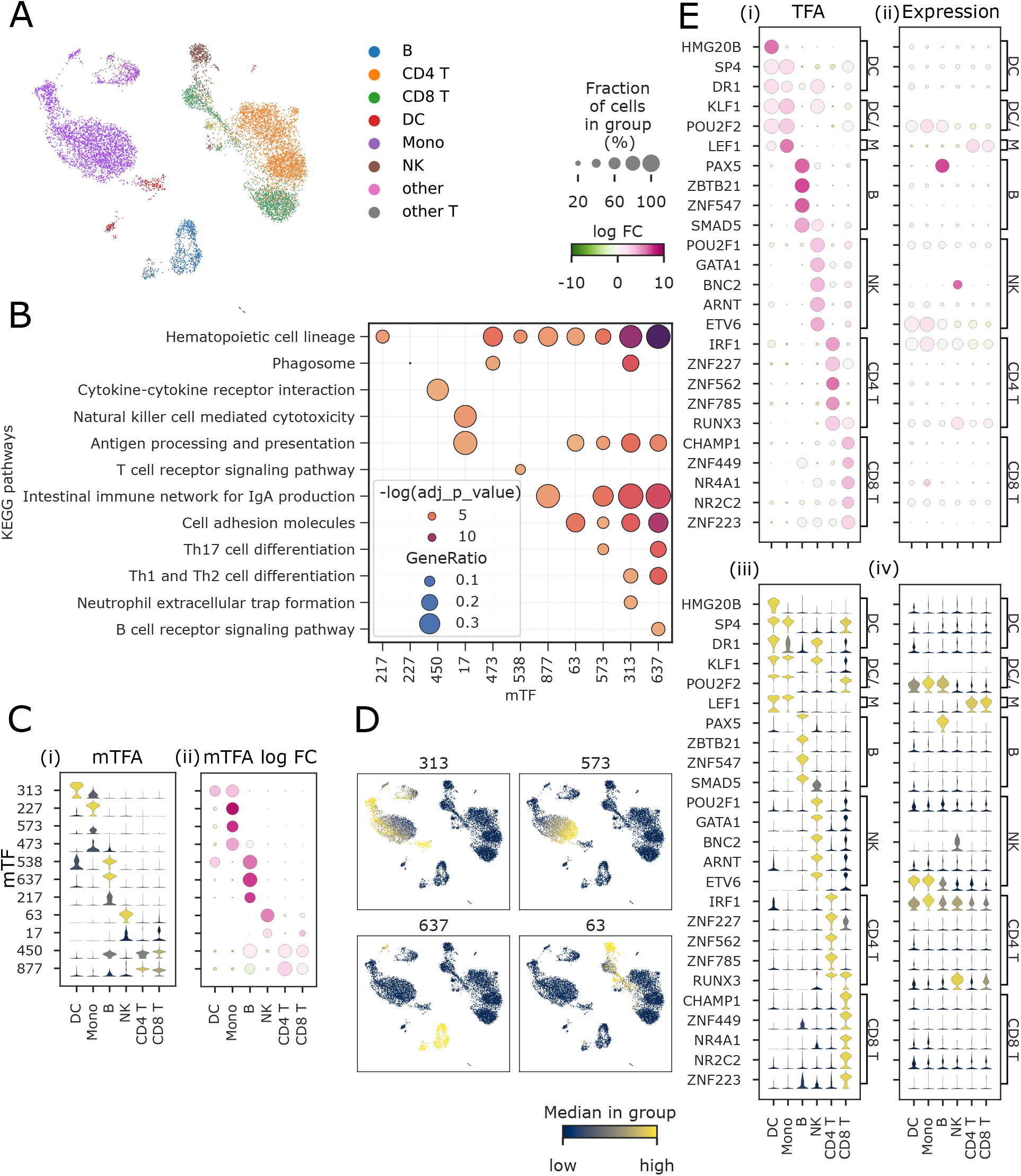
Single cell PBMC TFA and mTF functional enrichment analysis. TFA and mTFA are estimated from a JRL SupirFactor model. mTFs are associated with latent features 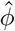 and are numbered 0-881. **A** UMAP projection of the single cell PBMC dataset, labeled with cell type annotations. **B** Selected enriched terms and associated mTFs for cell-type specific KEGG pathway based on mTF target genes, *ξ*^2^ > 0.01. **C** Activity of functionally enriched mTFs, over each cell-type. **D** UMAP projection of PBMCs colored by mTF activation. **E** Significant TF activation for specific cell-type populations and corresponding gene expression (scaled [0, 1] for comparison). Dot plots of (i) TFA and (ii) TF mRNA expression. Violin plots of (iii) TFA and (iv) TF mRNA expression.

PBMCs are a heterogeneous pool of multiple cell types, and each cell type may have subsets of cells in different states. We annotate these cells as dendritic cells (DC), monocytes (Mono), natural killer cells (NK), B-cells (B), CD4^+^ T-cells (CD4 T) and CD8^+^ T-cells (CD8 T). To account for this heterogeneity, we build a context aware Joint Representation Learning (JRL) SupirFactor model (Section 4.2). This allows the model to weight regulatory evidence based on context and aggregate into a joint latent feature space to build a joint SupirFactor GRN model. We define cell contexts for JRL by clustering cells using Leiden clustering[49], with a resolution = 0.2, generating 7 clusters (Supplemental Figure S11A-B). The PBMC SupirFactor model was trained using JRL on ATAC masked data (epochs=400) and subset to the most explanatory model weights once (epochs=100), resulting in a model that is scored on held-out cells (*R*^2^ = 0.35). The modeled PBMC regulatory network predicts 818 active TFs regulating 13698 genes, connected through 492 mTFs in a latent mTF layer (*ξ*^2^ ≥ 0) (Supplemental Figure S11C).

Eleven of these mTFs are linked to target genes which are functionally enriched for immune cell specific KEGG pathways (Figure 5B), and we interpret these mTFs as regulatory pathways. As an example, mTF-637 is active in B cells but largely inactive in other PBMCs (Figure 5B-D); the target genes of this mTF are functionally enriched for B-cell receptor signaling (Figure 5B). mTF 313 is activated in dendritic cells (DCs) and regulates genes functionally enriched for phagocytic activity and for Neutrophil extracellular trap formation, which activate DCs and allow them to mature [50]. mTF 227 is also linked to phagocytosis, although is principally active in monocytes. mTF 17 is active in NK cells with functional enrichment for Natural Killer cell mediated cytotoxicity genes. Overall, this demonstrates the utility of SupirFactor mTFs as a tool for identifying cell type specific regulation.

We can further examine TFs that have inferred cell-type specific activity (Figure 5E). SupirFactor distinguishes activation between myeloid (monocytes and DCs) and lymphocytic (B, NK and T cells) lineages (Figure 5E). The framework also recovers known cell-type specific regulators along the myeloid lineage, monocytes and DCs, (KLF1) [51], B cells (PAX5, SMAD5) [52, 53], NK cells (ARNT) [54], CD4 T cells (IRF1, RUNX3) [55, 56], and CD4 T cells (NR4A1) [57]. We show that SupirFactor infers cell-type-specific differential mTF activation and TF activation among distinct cell types that correspond with known biological processes and protein activity for multiple key cell types. The analysis of SupirFactor performance on the PBMC dataset demonstrates that SupirFactor can learn biologically relevant interactions in complex organisms and datasets.

## 3 Discussion

In this study we describe *StrUcture Primed Inference of Regulation using latent Factor ACTivity* (Supir-Factor), a model within the class of knowledge primed deep learning models. SupirFactor explicitly treats transcription factor activity as an interpretable latent state which drives gene transcription. This model uses a single objective function where the influence of the prior regulatory structure is optimized together with the GRN. SupirFactor combines the power of DNN optimization with prior structure constraints for inferring GRNs and explicit estimation of TFA. These TFA estimates are bounded by a ReLU activation function, and are directly quantifiable and interpretable on a per-observation basis.

SupirFactor has been carefully benchmarked using both bulk RNA expression data sets and single cell RNA expression data sets. We rely on model organisms *B. subtilis* and *S. cerevisiae* for benchmarking, as these organisms are well-characterized and have a partial experimentally validated ground truth network available, which we use for scoring recovery of GRN structure. This model organism benchmarking is important, as mouse and human data sets used for GRN inference benchmarking often lack reliable ground truth networks for scoring, and are restricted to using predictive metrics which have limited value. This benchmarking shows that the SupirFactor framework is versatile and has improved GRN inference over a comparable framework that relies on statistical learning, measured by recovery of network edges which are held out of the modeling. The SupirFactor GRN models are also predictive, and we expect that future work will tune the model to optimize network recovery using AUPR, or other model selection appropriate metrics, of held out gold standard network edges, by maximizing R^2^ for predictive power.

SupirFactor provides a novel metric for evaluating DNN architectures, specifically designed for the needs of GRN inference. Explained relative variance (ERV) estimates the importance of each latent feature to each output feature, directly and indirectly. This metric has appealing properties; ERV is bounded and facilitates ranking regulatory relationships and discarding non-predictive model weights. DNN models are often overparameterized, which results in problems interpreting model weights for GRN inference. As biological GRNs are sparse model selection must be part of the evaluation of any GRN inference algorithm. The model evaluation metric and model selection criteria we propose are also useful for evaluating the contribution of intermediate DNN layers which are not explicitly defined as TFs. By using ERV to evaluate linkages from regulatory TFs, through a latent meta-TF layer, to target genes, we are able to use the meta-TF layer as a powerful pathway analysis tool.

We demonstrate the validity of ERV by comparing it directly to GRN inference using model weights alone, and find that it improves GRN inference link interpertation. ERV also allows for post-training analysis of any gene expression data to determine which parts of the network are specific to that context. We show this by extracting context-specific networks from the *S. cerevisiae* single cell data set, which contains observations from fifteen different growth conditions. The recovery of these post-training context-specific networks is an improvement over previous work, which requires that the context is embedded into the model pre-training. SupirFactor is therefore a valuable tool to identify context specific and contextual regulatory interactions.

The driving features of GRNs can be condensed into TFs. Another core concept explored in this work is that of latent feature inference and interpertation, *i.e*. TF activity (TFA). In the model organisms *S. cerevisiae* we demonstrate that the model latent feature activity, the TFA of a TF is distinct from the expression of that same TF. We do this by studying the cell-cycle where we see a clear delay of relevant TFA compared to expression. Demonstrating that reliance on expression of TF as independant features to model a GRN does not capture the regulatory structure of GRNs. A drawback in many works related to GRN inference.

To demonstrate that SupirFactor scales to the complexity of mammalian systems we evaluate a model learned from a PBMC multi-ome single cell dataset, and characterize the pattern of TFA and functional enrichment in different contexts. The model make use of context specific prior evidence to further restrict TF variance. And we find that we can extract functional enrichment based on annotated celltypes reliably.

Reuse of computational models can be valuable as a tool to understand and conceptualize new experimental data evident by recent reuse of single cell sequencing atlases in the field of genomics[48, 58]. Unfortunately, the reuse of GRNs themselves is rare, and for most studies gene regulatory networks are inferred entirely based on new data. We consider this to be a general limitation in current-generation GRN inference models, which do not have mechanisms to embed new data into an existing GRN. SupirFactor tackles this by using a DNN architecture together with the transparency framework (ERV). We demonstrate this reuse by embedding novel data and by contextually analysing sub-networks after the model has been trained, gaining insight not explicitly provided to the model before training.

Developing explainable deep learning models for GRN inference is a critical requirement for improving models of gene expression and regulation[23]. The goal of this work was to build a formalized GRN inference model with explicit optimization and objective functions, from which latent states can be directly interpreted. The resulting formalism, SupirFactor, is a powerful GRN inference tool with additional pathway analysis and protein activity functionality, that can be applied to both bulk and single cell data. SupirFactor can harmonise regulatory evidence, epigenetic data and expression readout in a regulatory and functionally meaningful way. While challenges still exist, like model stability and model selection; tightly connected to the nature of non-linear machine learning algorithms, advances in single cell multi-omics and epigenetic sequencing are steadfast and will further narrow the specificity in model constraints with its inclusions.

With additional work related to architecture, algorithm development, and prior evidence construction, the framework can be further extended and prove even more useful.

## 4 Methods

### 4.1 *StrUcture Primed Inference of Regulation using latent Factor ACTivity* (SupirFactor)

We define two SupirFactor models. The shallow model, used mainly for testing, which consists of a single layer that represents individual TFs and their activity. The hierarchical model, which consists of two layers, the first representing individual TFs and their activity, and the second representing TFs aggregated into pathways, termed meta TFs (mTFs). The hierarchical model is the main model used in this work.

#### 4.1.1 Shallow SupirFactor

Starting from our model framework (Section 2.1), gene expression is a function of TFA and is used as the independent feature to weight influence on gene expression from TFs. We can formulate the problem as

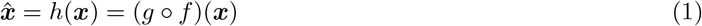

where 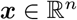 is the gene expression of one observation with *n* genes, with *g, f* as functions of the form

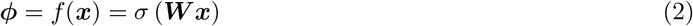

and

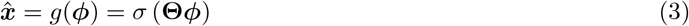

with the activation function *σ*, aggregating the linear combination of inputs for each latent feature. The linear combination of inputs without activation is similar to the NCA framework described in section 4.4.4 with the distinction that this formulation weighs ***W*** instead of fitting ***ϕ*** with a static ***P***.

**Θ** and ***W*** are the weighted connections of input features to output features in a single layer, with equation (3) corresponding to the shallow SupirFactor (Figure 1C). ***ϕ*** is the inferred latent activity interpreted as TFA of expression mapped through *f*. We set ***W*** ∈ ***P***, such that the sparsity structure of ***W*** is identical to ***P***. This ensures that (I) 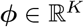, with *K* TFs, constraining the informative genes influences to those in the prior ***P***, and (II) the causal flow from regulator to target defined by the prior is enforced. Causality does not imply direct binding but rather in this case the variance of each TF being constrained to the variance of its targets as opposed to the covariance of the TF expression.

#### 4.1.2 Hierarchical SupirFactor

In Hierarchical SupirFactor (Figure 1A) we add an additional mTF layer between TFs and output gene expression. This allows higher order interactions between TFs (representing biological concepts like redundancy, competition, and physical complexing), and other conditional non-linear dependencies to be modelled. This extends the formulation of SupirFactor so that it can generate TF interaction hypotheses and be used as a tool for pathway analysis.

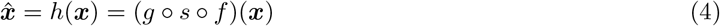

where

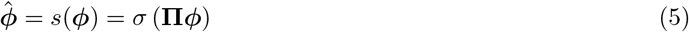

and

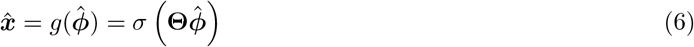

**Π** is the weight matrix of TF-TF interactions that maps individual TF activity to the mTF layer.

### 4.2 Joint representation learning for context specific constraints

Joint representation learning is a transfer learning method where context-specific evidence is aggregated into a common model structure [see 44, chap. 15] (Figure 1E). This is implemented in SupirFactor by adding a biological context-specific constraint on the prior evidence. We define ***P***_*C*_ as prior evidence for *C* where *C* is a biological context, like a cell type, growth condition, or temporal group. Weights ***W***_*C*_ ∈ ***P***_*C*_ are also context-specific, and are mapped jointly through **Π** and **Θ** that are common to all contexts. Experimental data is labeled with the appropriate context and data for each context is submitted batch-wise to the model for training. Context weights ***W***_*C*_ are individually trained and may vary between contexts if ***P***_*C*_ is the same. Equation (4) then takes the form;

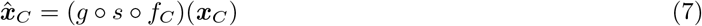

where *f_C_* is the context specific structure (***P***_*C*_) primed encoder.

#### 4.2.1 Fitting model

To train the SupirFactor model, ***W***, **Π**, and **Θ** are fit to minimize mean squared error between 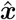 and *x*.

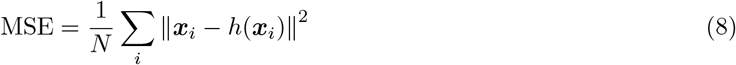

This is implemented as minimization by batch stochastic gradient descent with the Adam solver [59] and pytorch [60].

Nonzero encoder weights are initially mirrored and made non-negative in both the decoder **Θ** and encoder ***W***, although all elements in **Θ** are free to be fit by the solver. The prior assumption is that TFA is positively correlated with gene expression during model initialisation if no other evidence is available.

### 4.3 Model regularization

#### 4.3.1 Parameter penalties

Model weights are penalized, regularizing models to mitigate overfitting and to balance bias and variance [61]. We use a *weight decay* factor, corresponding to a ridge penalty [see 62, section. 6.7.6]. The objective function to be minimized is then

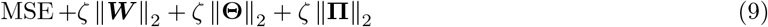

where *ζ* is the ridge penalty applied. The *ζ* parameter is set by cross validation, splitting the data into equal training and validation sets and evaluating model performance where *ζ* ∈ {0, 10^−10^, 10^−9^,…,10^−1^} [33, 63].

#### 4.3.2 Dropout

Dropout is an additional regularization method [64] where a fraction of nodes are removed from each sample during training, mitigating the risk that noise in the data will trap the model in a local minimum. Dropout can be applied to input or to latent layers. In short, training data is randomly batched into groups, and each batch is then used to train a network where a fraction, *p* of data points are removed randomly from each sample in each batch before feeding it through selected layer(s). This is implemented in SupirFactor through the Dropout module in pytorch. Droput is tested by cross validation as above on both the input and the latent TF layer, searching *p* ∈ {0, 0.01, 0.05, 0.1, 0.2, 0.3, 0.4, 0.5}.

### 4.4 Explained Relative Variance (ERV)

#### 4.4.1 Model selection and model parameter ranking

GRNs are sparse and most genes have a limited number of directly regulating TFs. The importance of model parameters is quantified, and relatively unimportant parameters are shrunk to zero. Two different ways of ranking inferred interactions are evaluated in this work: (I), Ranking the magnitude of model weights |*θ_i,k_*| (MODEL), and (II), ranking interactions by their explained relative error (ERV). ERV perturbs latent features and quantifies the consequence of that perturbation [37]. The goal is not to eliminate redundancy, but rather to eliminate over-parameterization and constrain the parameter space of SupirFactor.

ERV is calculated as coefficient of partial determination [43] so the bound on the error contributing to predict gene expression can be evaluated *ξ*^2^ ∈] −∞, 1] where a predictive link has an *ξ*^2^ ∈]0,1]. The GRN model is trained once, as re-training for each perturbed latent feature is computationally intractable. ERV is determined from the ratio of the full model MSE to the perturbed model MSE.

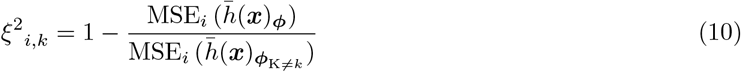

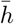 is the trained model, MSE_*i*_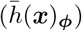 is the MSE of the full, unperturbed model for gene *i*, and MSE_*i*_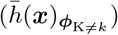 is the MSE of the perturbed model for gene *i* where the activation of the latent layer for regulator *k* is set to 0. Model parameters for each gene *i* are ranked by the value of *ξ*^2^_*i,k*_ for all *k*.

#### 4.4.2 Ranking model parameters for models with multiple layers

To be able to interpret any parameter in the model we use the ERV concept (section 4.4.1). To determine the importance in any hidden layer *L* not directly connected to the output we compute two *ξ*^2^ matrices. *ξ*^2^_*L*_ and *ξ*^2^_*L*+1_. Where *L* indicates the layer with corresponding latent feature input to the layer in question and *L* + 1 the next layer with corresponding latent feature input. For layer weights **Π** we compute each element *π_m,k_* with *m* as output feature and *k* as input feature by first computing the vectors *ξ*^2^_:, *k,L*_ and *ξ*^2^_:, *m, L*+1_. That is, the *ξ*^2^ of each element of **Θ**_:, *m*_ and the *ξ*^2^ of each element of the indirect contribution of latent feature *k* in layer *L* to the output genes *ξ*^2^_:, *k,L*_. To eliminate weights in **Π** we threshold ***ξ***^2^_:,*k,L*+1_ > *∊*_1_ and ***ξ***^2^_:,*m L*_ > *∊*_2_ and compute the ERV;

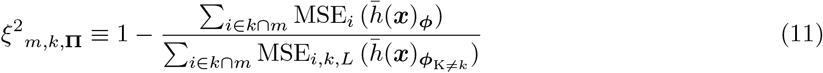

where *i* ∈ *k* ∩ *m* is the intersection set of predictive TF to gene interactions for latent feature *k* from layer *L*, and *m* for layer *L* + 1. If *k* ∩ *m* = θ then *ξ*^2^_*m, k*,Π_ ≡ 0.

The classical GRN realisation is interpreted as the indirect connections from *ξ*^2^_*i,k,L*_, connecting the latent input features in layer *L*, *k* to the output target genes *i*. With *L* in this case representing the TFA activation layer.

#### 4.4.3 Stable architecture and non predictive weight elimination

To select an interpretable model we want to reduce the model size in terms of individual weights to arrive at a model with parameters that are predictive and stable. We define predictive as when an individual parameter that can be determined to connect an upstream regulator to its downstream target, has *ξ*^2^ > *∊*. Stable in this case means that the parameter is predictive on unseen data when the model is trained on reduced subset of parameters.

We apply model constraints on individual weights. Parameter weights 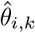 (and *θ_i,m_* in hierarchical Supir-Factor), with *i* output and *k*, and *m* input features are removed if *ξ*^2^_*i,k,L*_ ≤ *∊* for the respective layer *L*, with a choice of *∊* = 0 *i.e*. not predictive of the output, is the most conservative.

To enforce these constraints in number of regulators per gene we use a model selection step after an initial training run (Figure 1E). The model selection step is derived from *ξ*^2^ where

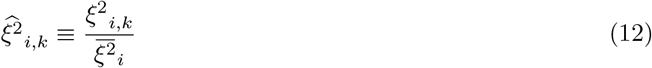

with 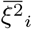 as the maximum *ξ*^2^ for gene *i*. Selection is done iteratively by selecting a threshold *∊*. The model is then refit with parameters where 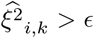. This is done iteratively until convergence where at each step *ξ*^2^ is recomputed. *∊* is a measure of inclusion of relative predictive power. Using *∊* = 0 means all predictive links are kept after sub-setting and relative predictive power 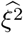 is not impacting the subset.

Using 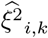 facilitate selecting regulator subsets where, unless the gene is too noisy or the prior lack sufficient information that can reliably predict the specific expression pattern (*i.e*. all *ξ*^2^_*k*_ ≤ *∊* for the gene in question), at least 1 TF can be inferred to be predictive relative to all *k* for that gene *i* and other regulators are ranked relative to it.

For the hierarchical model sub-setting, weights are eliminated if no predictive interactions can be derived from the indirect path between a latent feature *k* through the latent feature *m* in the subsequent layer to the set of joint output features, above the chosen *e* threshold. If *ξ*^2^_*k,m*,Π_ ≤ 0 the hidden layer weight is pruned.

#### 4.4.4 Network Component Analysis (NCA)

Network component analysis (NCA) can be used to estimate TFA directly by formulating the causal network inference problem so that

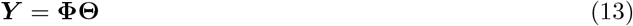

where 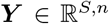 is gene expression with *n* genes and *S* samples, 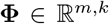 is the TFA with *k* TFs, and 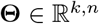 is the regulatory effects linking genes to TFs. The unknown “true” **Θ** is the regulatory interaction between genes and TFs we want to find. **Φ** is unknown given the assumption that the expression level of a transcription factor *k* does not correlate well with the activity of the protein [65]. Therefore we need to solve for both **Θ** and **Φ**, which forces us to convolve our estimation of regulatory effect and the TFA. To deconvolve and solve this we impose a prior ***P*** with elements ∈ {0,1} as an initial guess to the structure of **Θ** and use that to solve for an initial estimate for **Φ**.

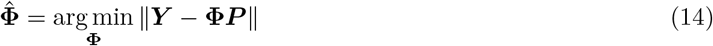

This is solved by ordinary least squares. The estimated **Φ** is then used to solve for **Θ**

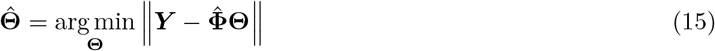

This is interpreted as an estimate of TFA given a number of reporter genes defined by the prior ***P***, *i.e*. the expression level of the target genes is a proxy for how active a TF is in any given sample. The variance of the TF activity is defined and constrained by the variance of the reporter genes.

### 4.5 Epigenetic masking

To incorporate the paired ATAC- and RNA-sequencing data we create a masking scheme to mask input gene expression profiles with the ATAC so that expression input

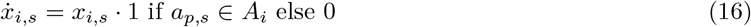

with *x_i,s_* being the expression of gene *i* in sample *s* and *a_p,s_* being an available ATAC peak *p* in the set of accessible regions *A_i_* associated with gene *i*.

### 4.6 4.6 Data Acquisition

#### 4.6.1 Bulk expression data

*Bacillus subtilis* bulk expression data for data set 1 (B1) [10] and data set 2 (B2) [66], and the known prior network [10] were used as previously described [41]. *Saccharomyces cerevisiae* bulk expression data for data set 1 [67] and data set 2 [68], and the known prior network [69] were used as previously described [41].

#### 4.6.2 Single cell expression data

*Saccharomyces cerevisiae* single cell training data was assembled from [13] and [70] as previously described [4].

*S. cerevisiae* test data was collected using a method previously published [13]. In short, biological replicates containing unique, transcriptionally-expressed molecular barcodes of a wild-type strain 1 (*MATα/MATa*Δ*ho::NatMX/*Δ*ho::KanMX*) and a wild-type strain 2 (*MATα/MATa HO/*Δ*ho::NatMX HAP1+::pACT1-Z3EV::NatMX/HAP1 ura3*Δ*0/URA3 can1*Δ*::prSTE2-HIS5/CAN1 HIS3/his3*Δ*1 LYP1/lyp1*Δ*0*) were generated as previously described [13].

Strains were grown overnight in rich media (YPD as previously described [13]) and then subcultured into 100mL YPD or minimal media (MMD as previously described [13]) for 3 hours. Cells from each flask were then taken, fixed with saturated ammonium sulfate, processed, and sequenced using the protocol as previously described [13]. Raw sequencing data was processed into count data using a previously-described pipeline [13] which joined the transcriptional barcodes to individual cells, assigning specific genotypes to cells and removing any cell containing multiple distinct barcodes as doublets. Four data sets were then created from this count table; YPD 1 (n=1531, wild-type strain 1 in rich YPD media), YPD 2 (n=1428, wild-type strain 2 in rich YPD media), MMD 1 (n=492, wild-type strain 1 in minimal MMD media), and MMD 2 (n=463, wild-type strain 2 in minimal MMD media). This data is deposited in NCBI GEO as GSE218089.

#### 4.6.3 Yeast Annotations

Cell cycle related yeast genes are annotated based on [71]. Ribosomal, ribosomal biogenesis, and induced environmental stress response genes are annotated based on [72].

Individual cells in single cell RNA expression data sets are assigned a cell cycle phase based on cell cycle gene annotations. Expression of each cell cycle gene is normalized to a mean of 0 and unit variance. All marker genes annotated with a specific cell cycle phase (G1, S, G2, M, or M/G1) are grouped, and the cell is assigned to the phase that has the maximum mean group expression.

Kyoto Encyclopedia of Genes and Genomes (KEGG) annotations [47] were selected to cover the majority of the core yeast metabolism. KEGG annotations are KEGG:04111 (Cell cycle), KEGG:00010 (Glycolysis), KEGG:00020 (TCA cycle), KEGG:00500 (Starches), KEGG:00660 (C5-Branched), KEGG:01210 (C5-Branched), KEGG:00250 (AA (A, D, E, N, Q)), KEGG:00260 (AA (G, S, T)), KEGG:00330 (AA (R & P)), KEGG:00400 (AA (F, Y, W)), KEGG:00270 (AA (C & M)), KEGG:00920 (Sulfur), KEGG:00061 (Fatty acids), KEGG:00230 (Purines), and KEGG:00240 (Pyrimidines)

#### 4.6.4 PBMC multi-ome dataset prepocessing

Paired PBMC scRNA-seq and scATAC-seq was downloaded from the 10x website (https://support.10xgenomics.com/single-cell-multiome-atac-gex/datasets/1.0.0/pbmc_granulocyte_sorted_10k). This data was preprocessed using a previously published workflow [48]. In short, the RNA-seq data is preprocessed as detailed in Section 4.7.1, with the additional filtering of cells with > 25000 or < 1000 counts and < 20% mitocondrial counts of total. For the ATACseq data we used epiScanpy[73], filtereing peaks in < 10 cells and cells with < 5000 or *>* 7 · 10^4^ counts, and with a variability score < 0.515. Final data contains 10411 cells, 21601 genes and 75111 peaks,

Cell types are annotated using the reference PBMC dataset[58] passed to scanpy’s [74] inject label transfer function, resulting in 8 annotated celltypes (Figure 5A).

#### 4.6.5 ENCODE PBMC prior knowledge network construction

TF-ChIP peaks were obtained as narrowPeak BED files from the ENCODE project database. The GRCh38 genomic annotations (NCBI GCF_000001405.39) were obtained as a GTF file from NCBI and filtered for protein-coding genes.

TF-ChIP peaks were linked to candidate target genes with the inferelator-prior package [41]. TF peaks were annotated as possible regulators of a gene if they were within 50kbp upstream of a gene transcription start site and 2kbp downstream of a gene transcription site, with no other gene between the TF peak and the gene transcription start site. TF Peaks were further filtered to remove any regulators not annotated as TFs (including GTFs, chromatin modifiers, and polymerase subunits).

This large pool of potential TF regulatory peaks were then subset for intersetion with annotated regulatory regions for PBMC cell types (ENCODE Accession IDs ENCFF776AJJ, ENCFF497NXM, ENCFF984SPH, ENCFF079TQT, ENCFF862ULW, ENCFF504FDC, and ENCFF905BHJ). The peak intensity (signalValue) was summed for all peaks annotated to each TF-gene pair to generate a genes by TFs putative regulatory matrix. This matrix was further constrained for sparsity by retaining at most 1.5% non-zero values for each TF, shrinking all values below this threshold to zero, and producing a genes by TFs prior knowledge network matrix with a sparsity of 1.22%.

### 4.7 Data preprocessing and model parameterisation

#### 4.7.1 Single cell pre-processing

For the single cell data, unless otherwise stated, we follow standard normalization procedures which include, (i) filtering genes with expression in < 10 cells, (ii) count normalization; scaling each cells total count to the same value over the dataset. This serves to eliminate the effect of variable sequencing depth in the experimental technique, and (iii) log transforming the (data + 1) using the natural logarithm.

#### 4.7.2 Feature normalization

For bulk data we use the standard normalization of each input feature so that

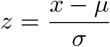

for each gene with *μ* = mean and *σ* = standard deviation over the gene.

To preserve the sparse structure of single cell data we, in addition to the above, adopt a robust normalization approach without centering. Each gene is scaled by the range of the 1 and 99 percentile and shifted so the lowest value for each gene = 0 implemented using the scikit-learn RobustScaler method [75] which we call RobustMinScaler.

### 4.8 ReLU in hierarchical SupirFactor

For DNN linear activation does not contribute meaningfully in different layers and can be reduced to a single linear map. The rectified linear unit (ReLU)[44] truncates activation to stay strictly positive and injects non-linearities into the model architecture. For hierarchical SupirFactor we therefore use ReLU and define the gradient for the ReLU function

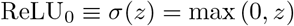

so that with *z* = 0 the gradient is ≡ 1. With *z* as the linear combination of inputs to each feature.

### 4.9 Visualisation

Visualisations throughout this work, if not stated otherwise, was generated in Matplotlib[76] with some components done with seaborn[77] and scanpy[74].

## 5 Acknowledgement

We thank past and present members of the Christiaen, Bonneau, and Gresham labs for discussions and valuable feedback on this manuscript. This work was supported in part through the NYU’s IT High Performance Computing resources, services, and staff expertise.

## 6 Funding

This work was supported by the NIH (R01HD096770, RM1HG011014 R01NS116350, R01NS118183, R01GM134066, R01GM107466), and the Simons Foundation.

---

**S1:**
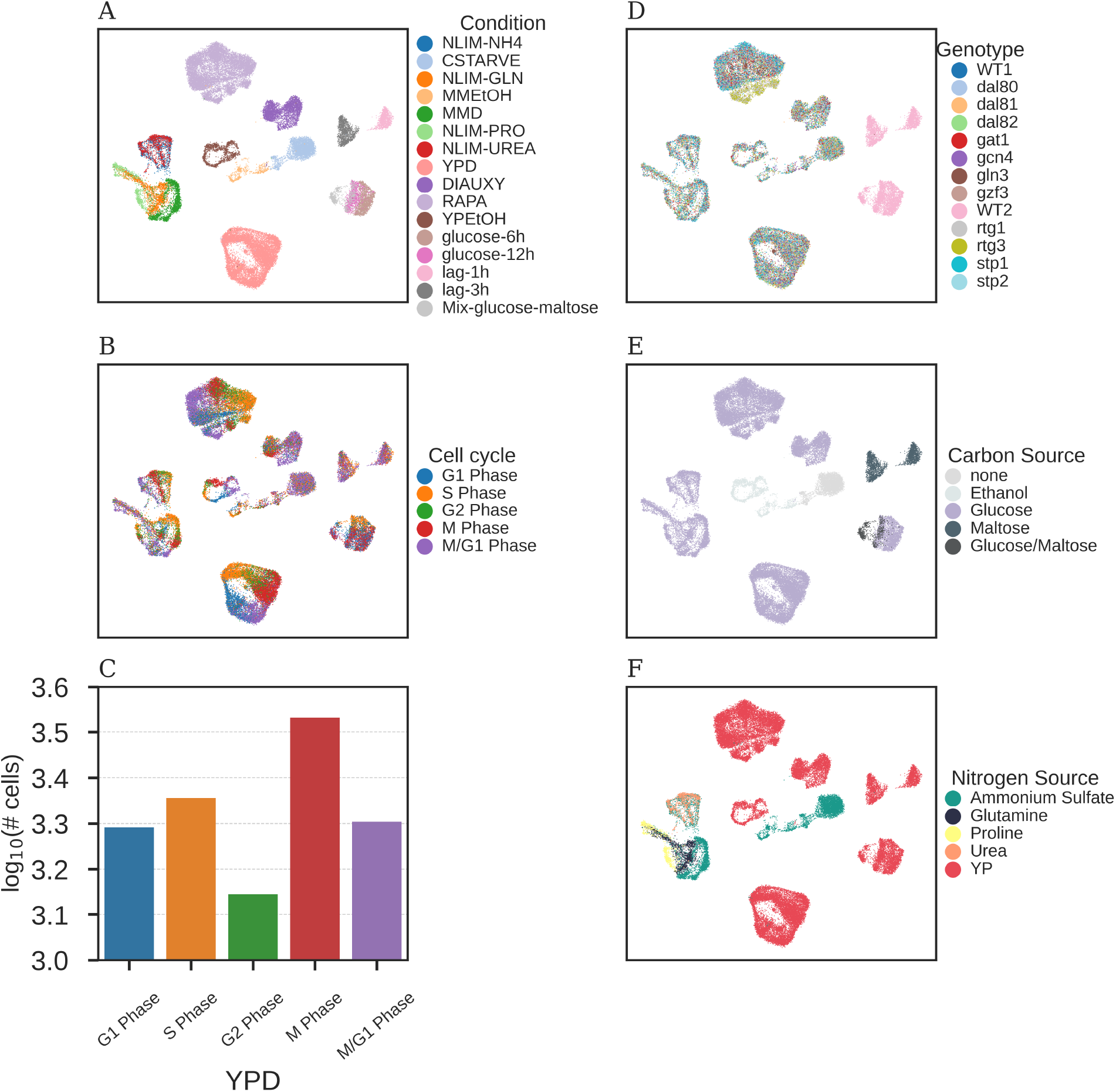
Overview of the single cell combined datasets [13] and [78] (scY) in *S. cerevisiae*. UMAP [79] projected data with annotation of **A**: growth condition, **B**: inferred cell cycle phase, and **C**: Frequency of cell cycle annotated cells in YPD. **D**: Genotypes. **E**: Carbon source for growth condition. **F**: Nitrogen source for growth condition.

**S2:**
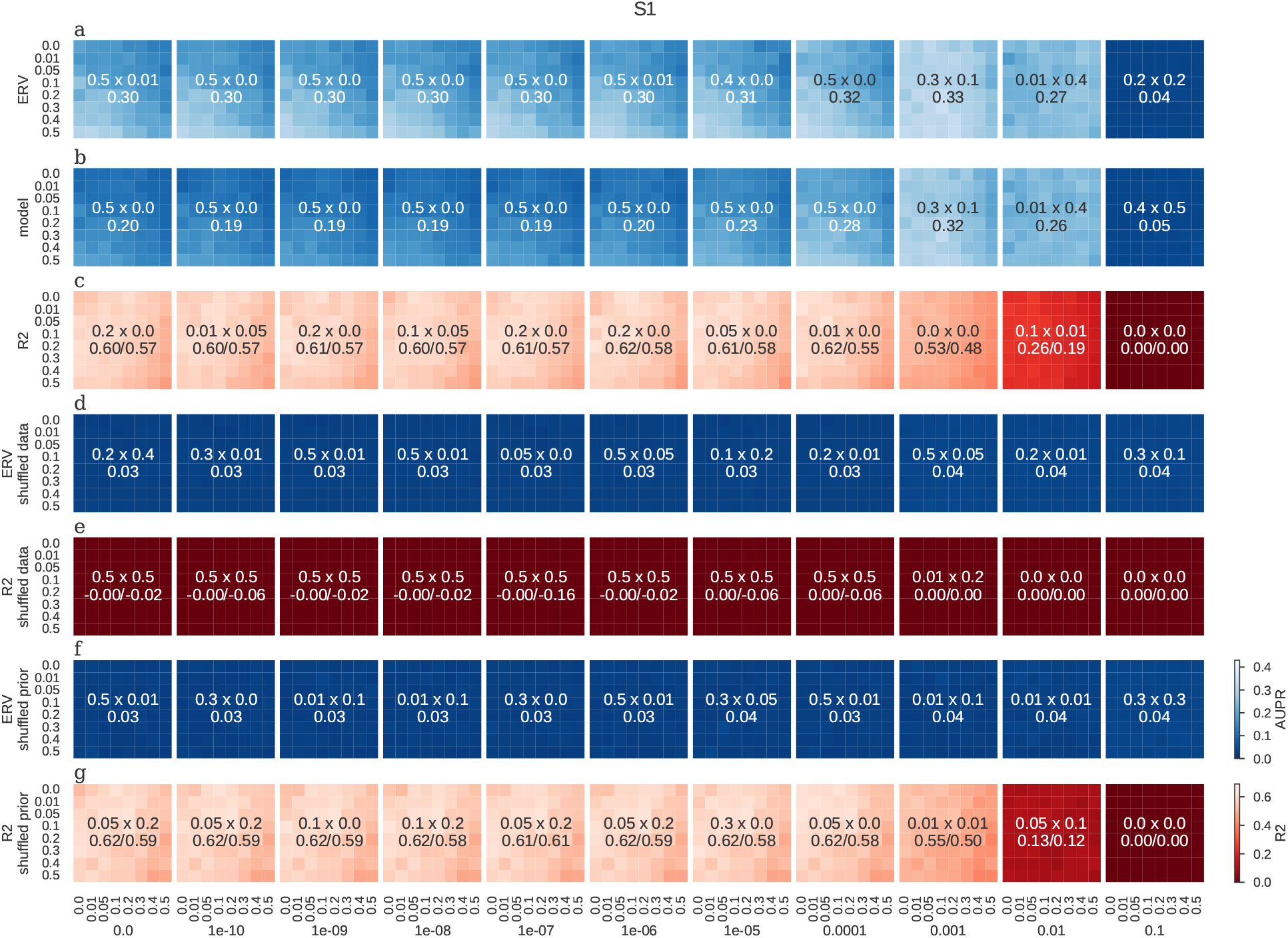
Dataset S1[68]. Hyper-parameter cube search. Each column of squares indicate a weight-decay (L2) penalty, with 0 and then 10 log linear values between 10^−10^ − 10^−1^. Each square is the grid search of dropout on input feature (y-axis) and latent features (x-axis). Rows in blue shade (**a**, **b**, **d** and **f**) indicate GRN recovery performance measured by area under precision recall curve (AUPR) on 20% held out of gold-standard network. ERV (**a**, **d** and **f**) indicate the GRN is ranking based on explained relative variance (Section 4.4.1), model (**b**) indicate network interactions are ranked based on their weights magnitude, with larger magnitude indicate more influential. Red shades (**c**, **e** and **g**) indicate prediction accuracy *R*^2^ on held out, n=50%, of samples. Negative controls are done by shuffling either the data (**d** and **e**) or the prior network structure (**f** and **g**).

**S3:**
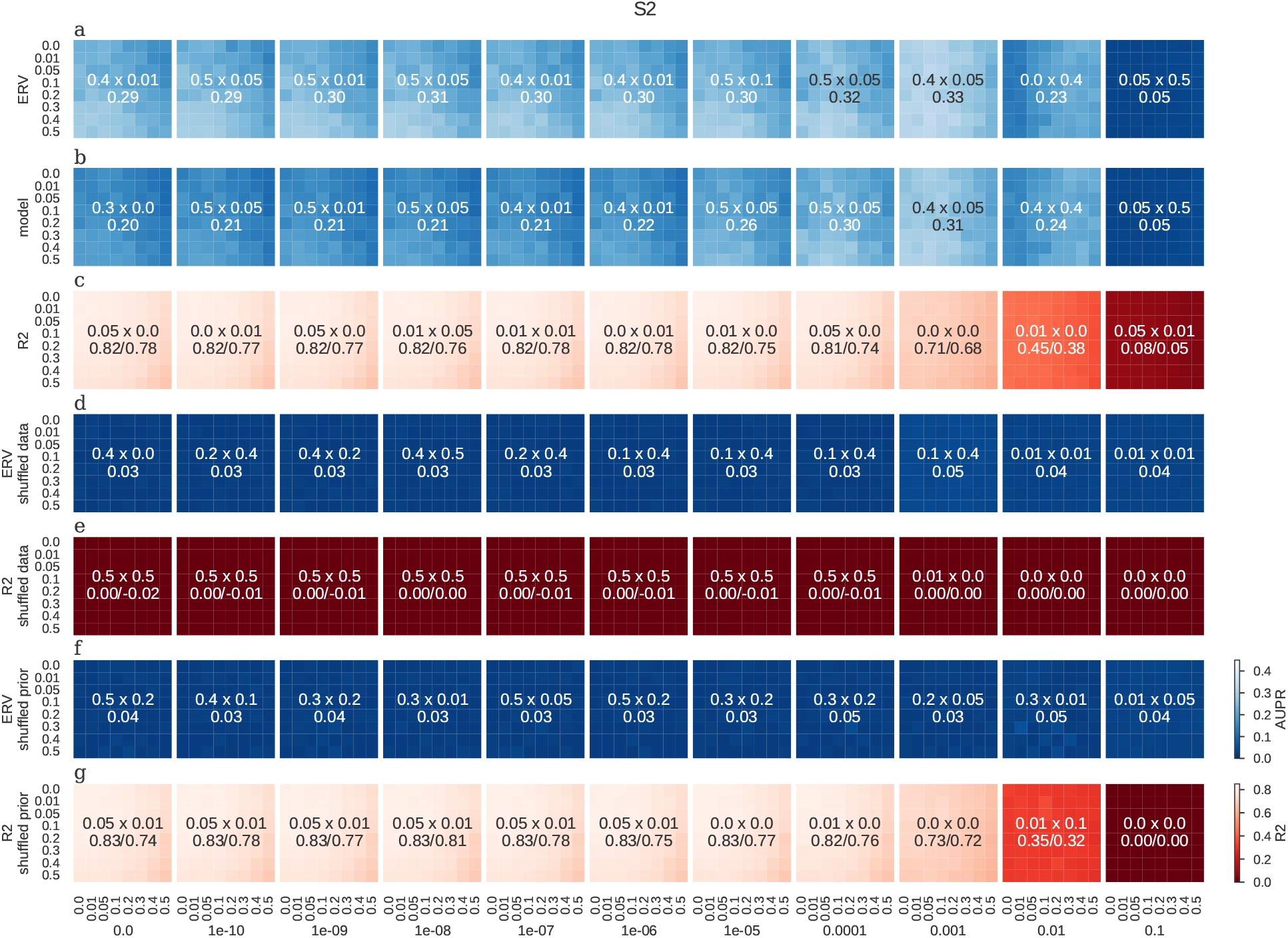
Dataset S2[67]. Description as in Figure S2

**S4:**
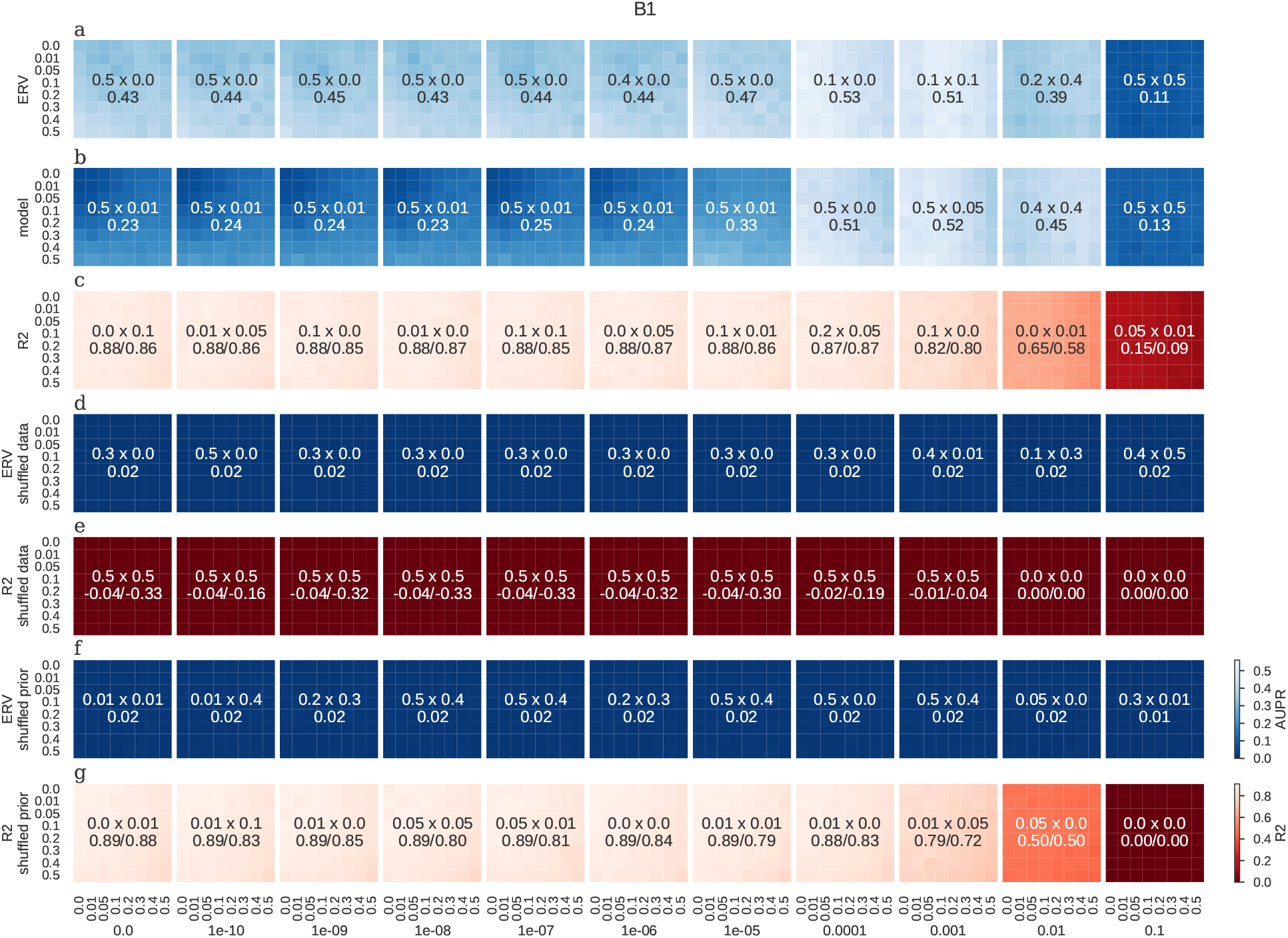
Dataset B1[10]. Description as in Figure S2

**S5:**
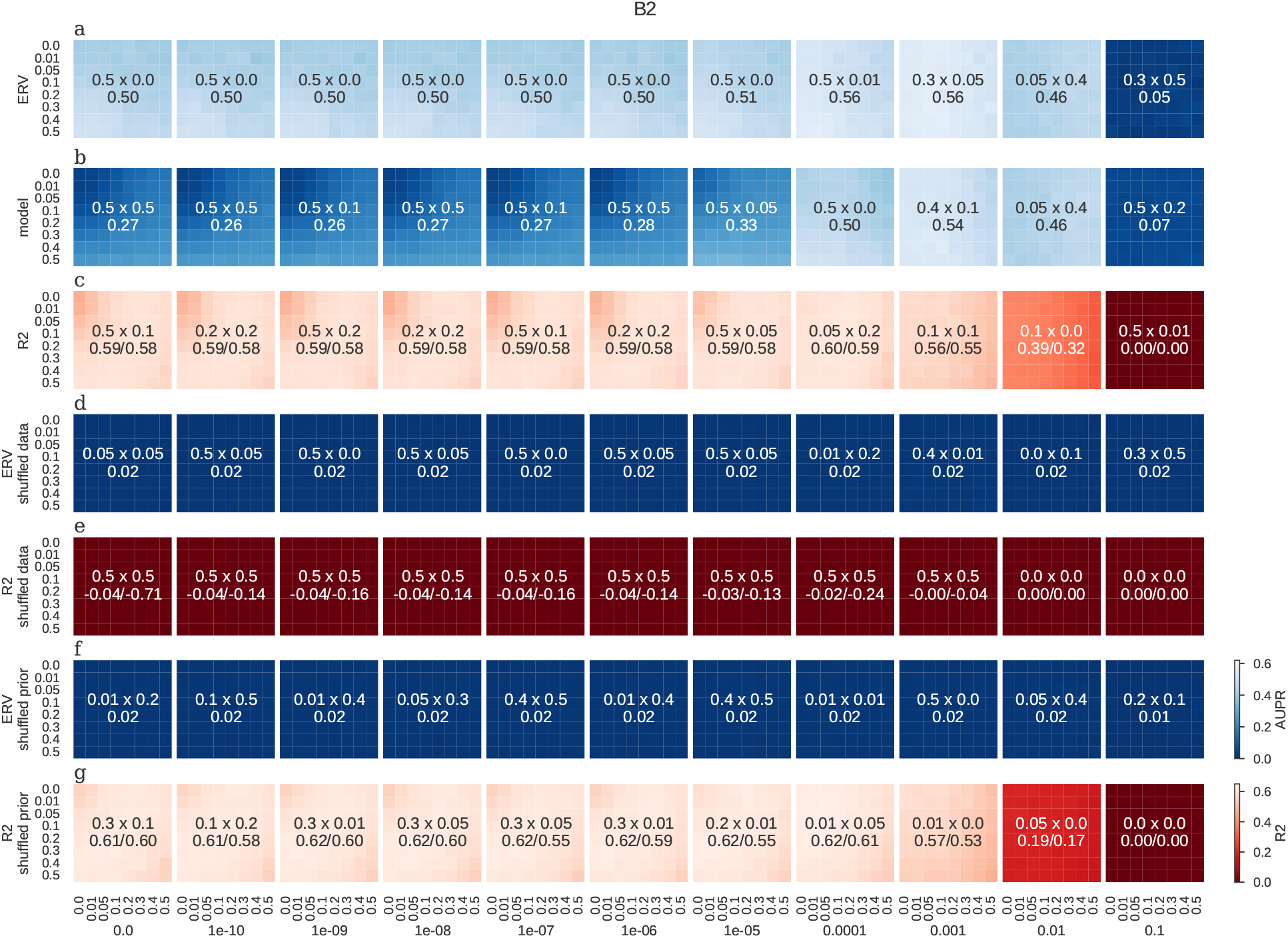
Dataset B2[66]. Description as in Figure S2

**S6:**
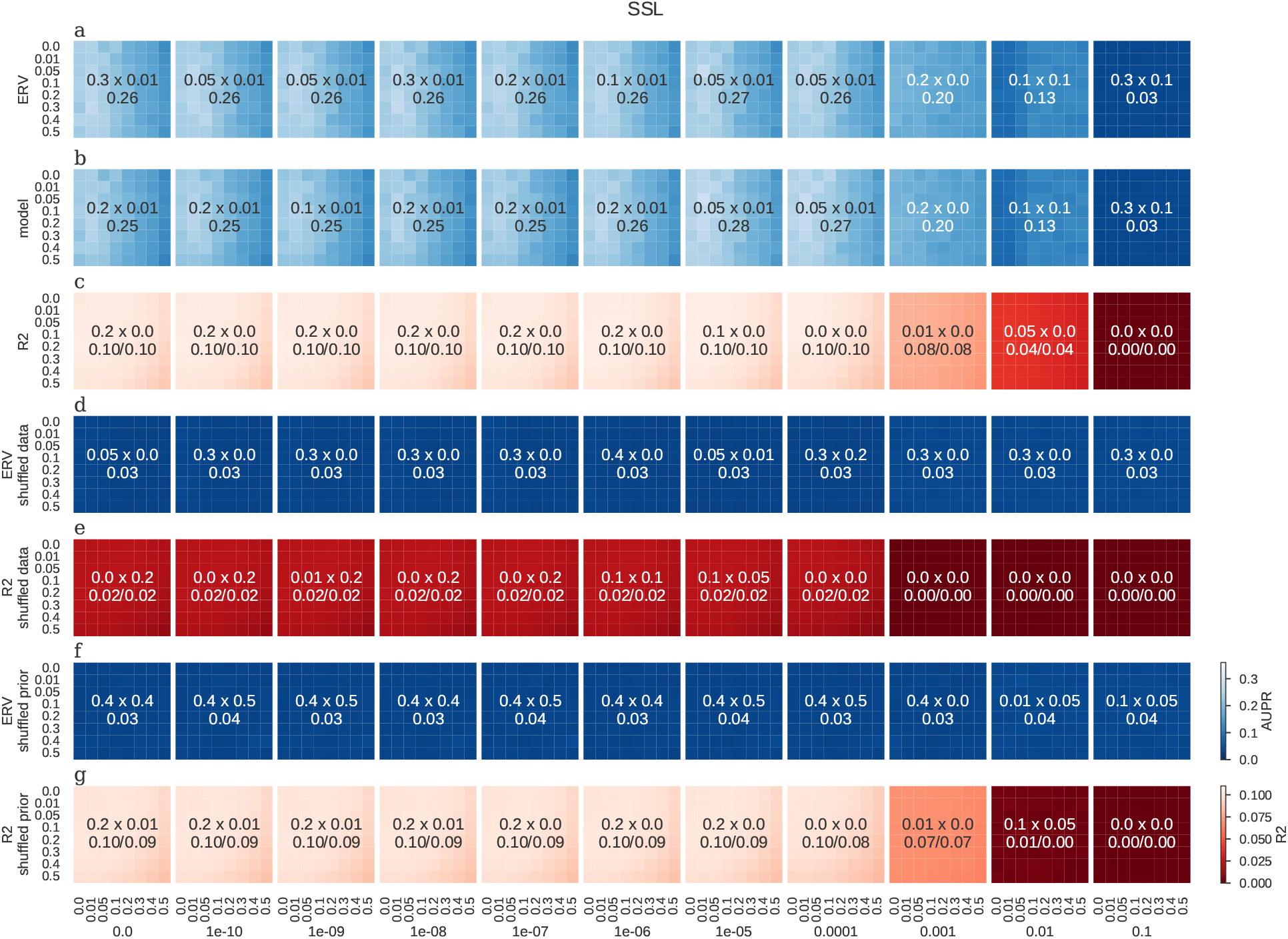
Dataset scY[13, 78] with StandardScaler normalisation and linear activation (SSL) configuration. Description as in Figure S2

**S7:**
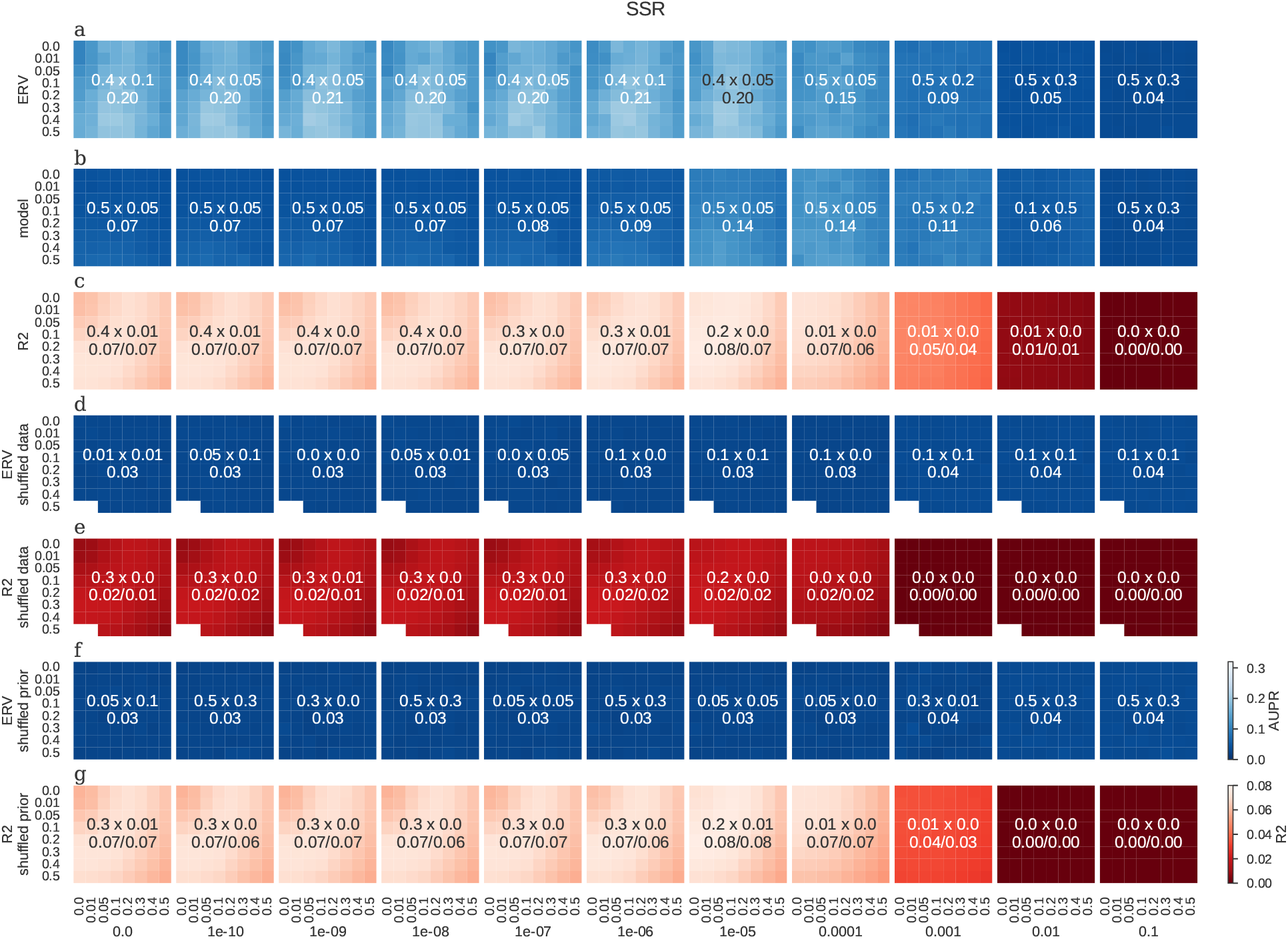
Dataset scY[13, 78] with StandardScaler normalisation ReLU_0_ activation (SSR). Description as in Figure S2

**S8:**
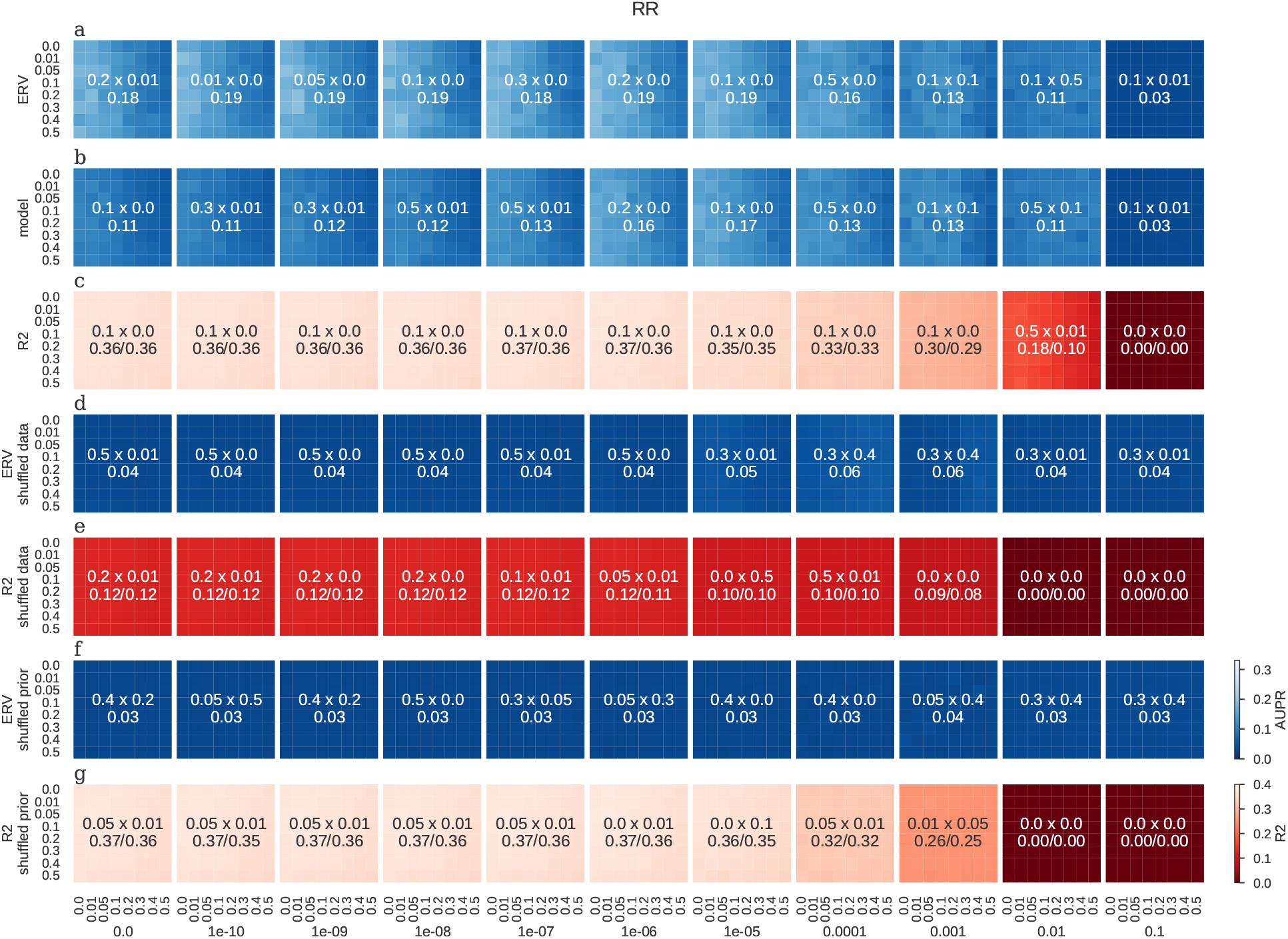
Dataset scY[13, 78] with RobustMinScaler normalisation and ReLU_0_ activation (RR). Description as in Figure S2

**S9:**
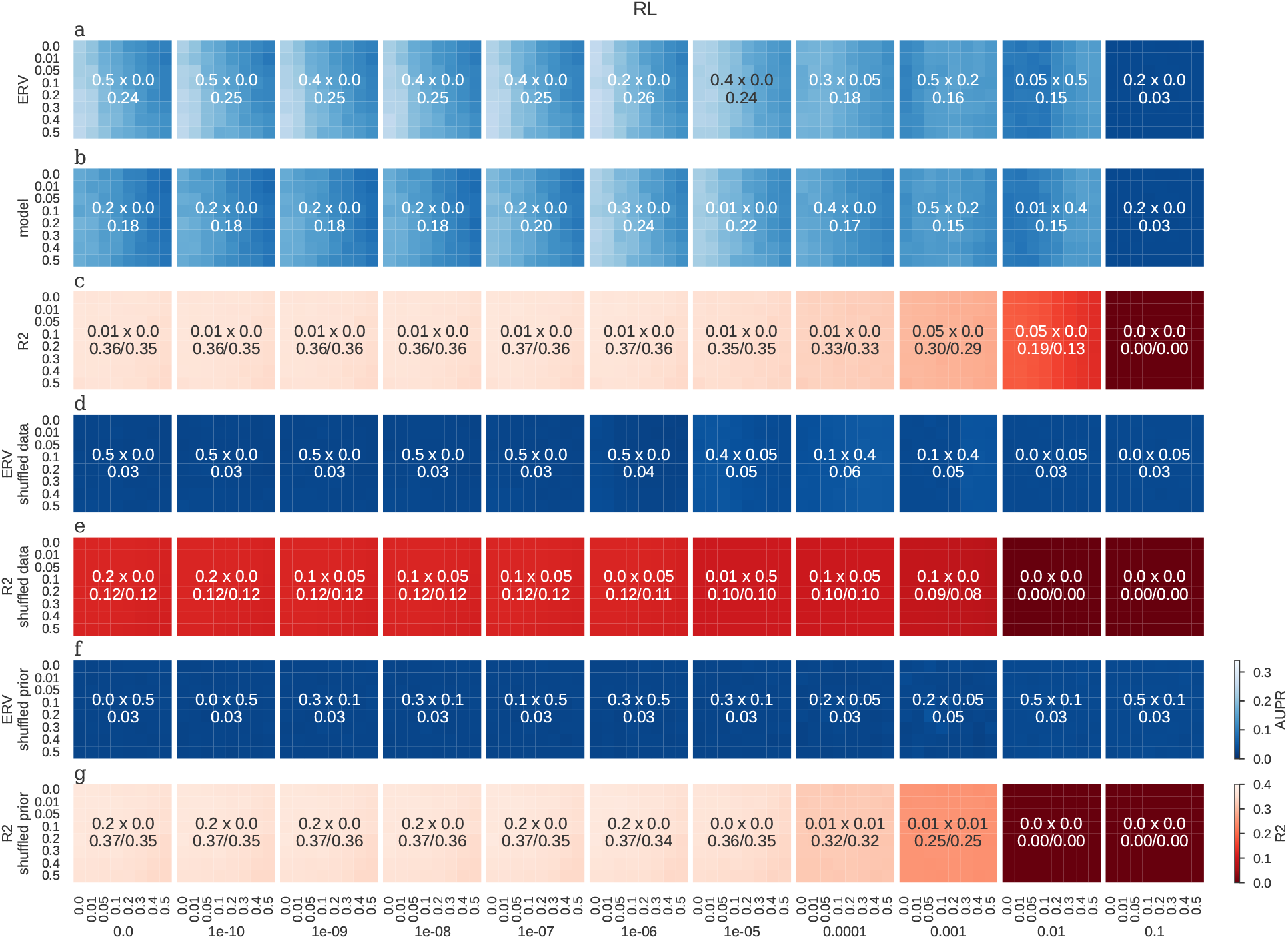
Dataset scY[13, 78] with RobustMinScaler normalisation and linear activation (RL). Description as in Figure S2

**S10:**
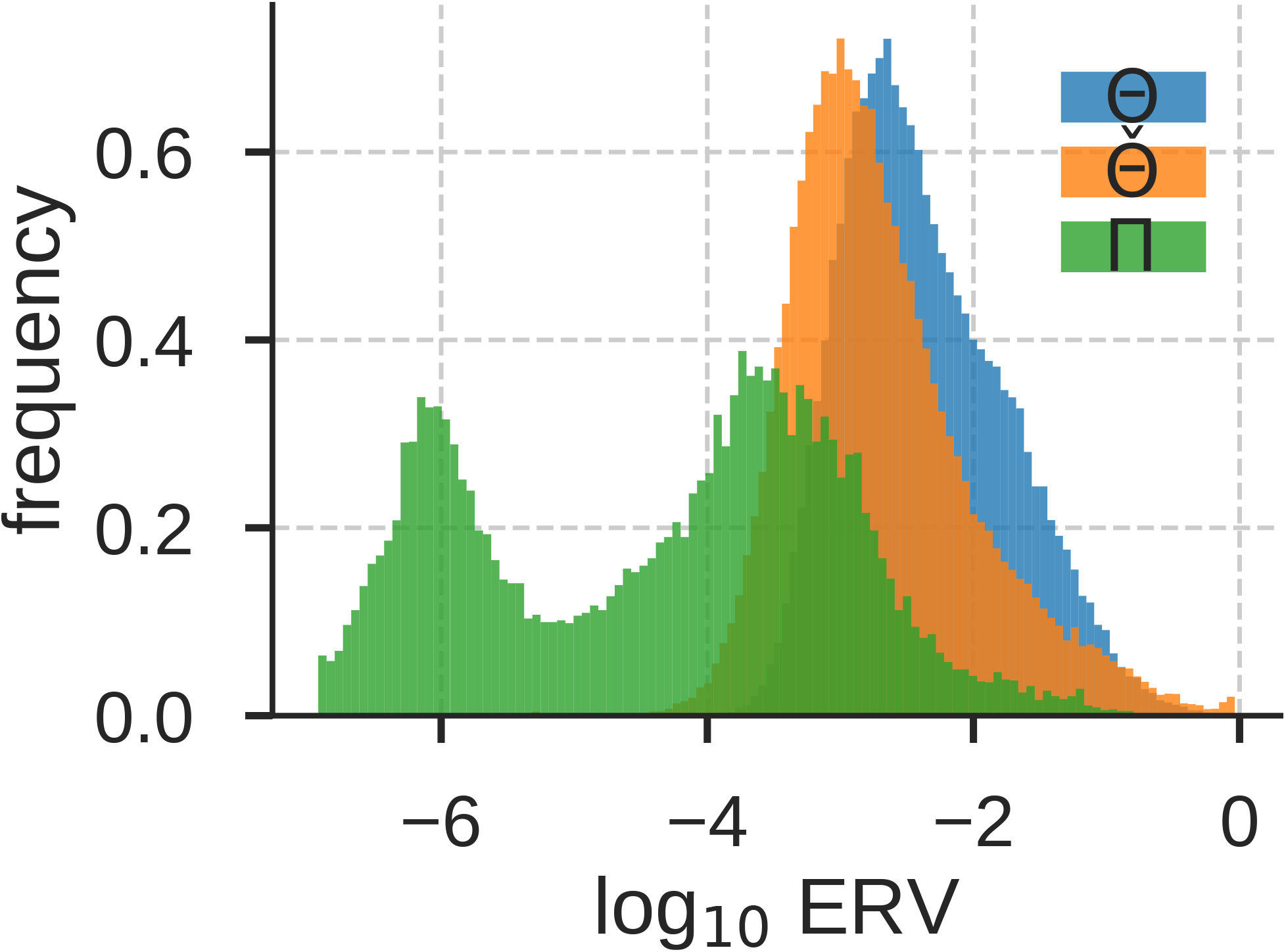
Distribution of ERV (*ξ*^2^) for hierarchical SupirFactor trained on scY.

**S11:**
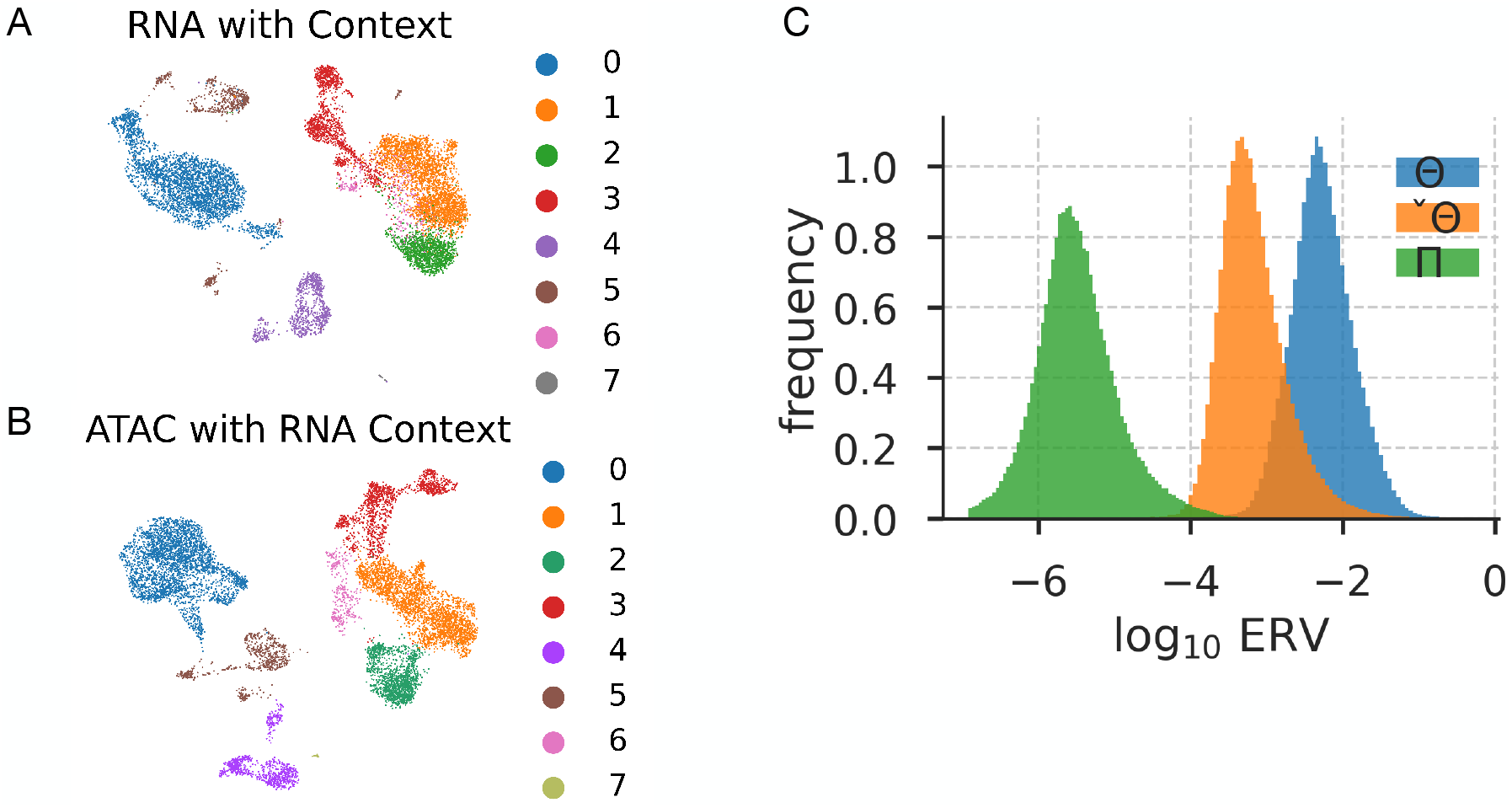
Single cell PBMC Contexts and Networks **A**: UMAP projection of the scRNA-Seq PBMC dataset with “Context” via leiden clustering (0.2) **B**: UMAP projection of the scATAC-Seq PBMC dataset with “Context” via leiden clustering (0.2) **C**: Distribution of ERV for 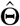, **Π** and **Θ** networks

**S12:**
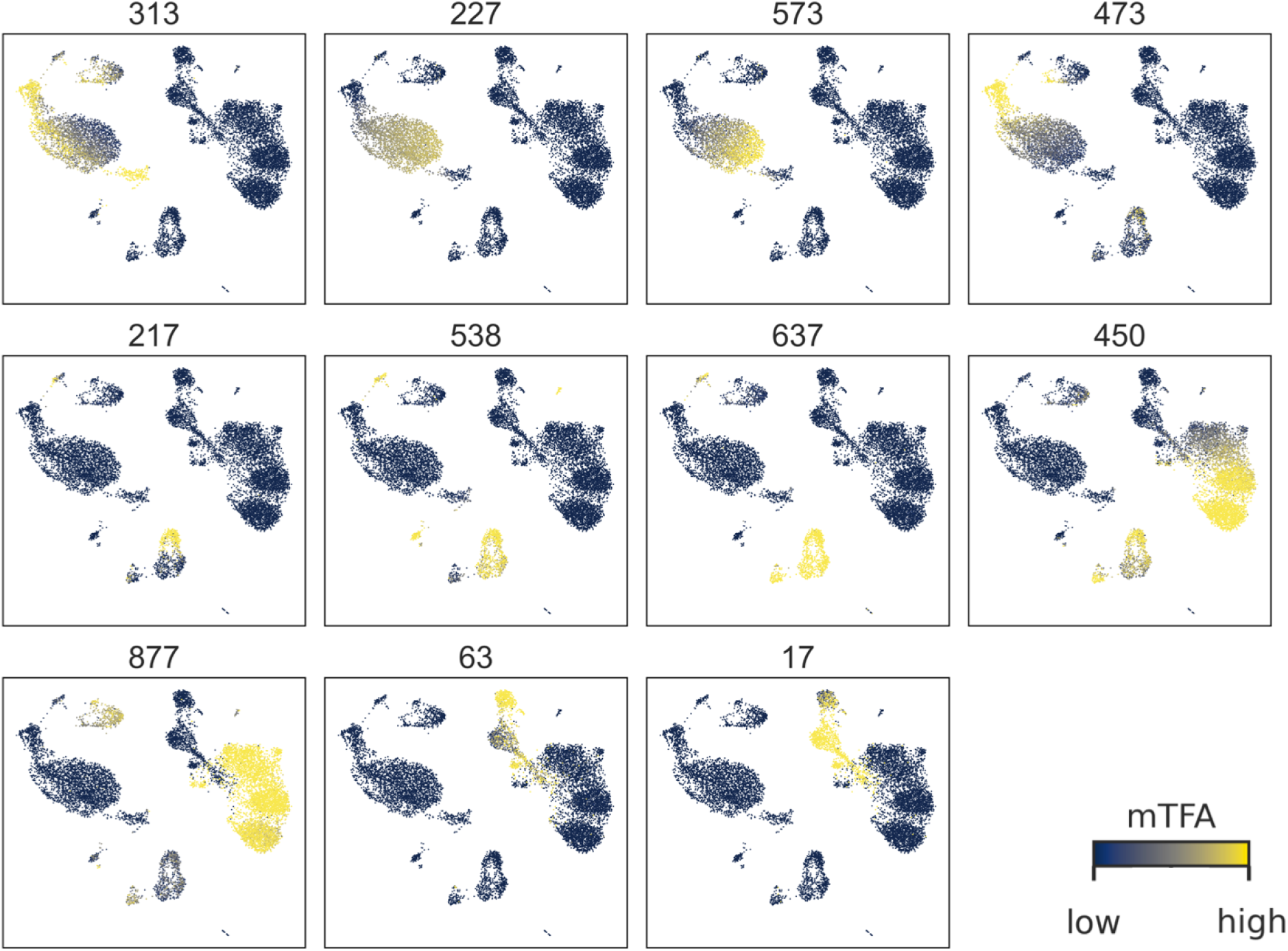
UMAP projection of all functionally enriched mTF activation (mTF activity scaled [0, 1] for comparison).

## Notes

### Competing Interest Statement

The authors have declared no competing interest.

